# Beyond Arrhenius: Nonlinear and negative temperature scaling of biological rates from multi-step mechanisms

**DOI:** 10.1101/2025.09.01.673554

**Authors:** Simen Jacobs, Federico Vazquez, Nikita Frolov, Lendert Gelens

## Abstract

Temperature shapes all biological processes, particularly during the early development of ectothermic organisms. A widely used framework for describing temperature dependence is the Arrhenius equation, which predicts an exponential increase in rates with temperature. However, biological rates often deviate from this prediction when measured across broader temperature ranges. While negative apparent activation energies are often attributed to protein denaturation, this cannot explain similar behavior observed at temperatures where enzymes remain stable. These broader scaling patterns remain mechanistically unexplained. Here we present a general Markov chain framework for modeling biological timing as cascades of reversible, temperature-dependent steps. Applying this model to 121 published datasets spanning diverse species, biological timescales, and temperature ranges, we find that a consistent three-zone scaling pattern emerges: Arrhenius-like behavior at low and high temperatures, and a quadratic exponential regime at intermediate temperatures. We show that this pattern arises naturally from differences in activation energies across steps in the network. The quadratic exponential regime is an emergent feature of averaging across many steps and is robust to variation across network realizations. In contrast, Arrhenius-like scaling at the extremes tends to be more variable and originates from smaller sub-networks. Apparent negative activation energies can emerge naturally from the dynamics of multi-step networks, even in the absence of protein denaturation. Our framework provides a unified mechanistic explanation for diverse temperature-scaling behaviors in biology and may help predict how developmental and physiological processes respond to environmental change. Although we focus on development, the model is broadly applicable to biological systems governed by multi-step reaction networks.

## Introduction

Temperature influences all living organisms, shaping their physiology, behavior, and overall fitness. This is particularly critical during early embryonic development, where temperature acts as a key regulator of developmental rates, impacting biochemical reactions, metabolic activities, and organismal performance. While endothermic organisms maintain stable internal temperatures, ectotherms are especially vulnerable to environmental temperature changes, making them sensitive to the effects of climate change. Understanding how temperature governs biological processes is thus essential for predicting the consequences of global warming on biodiversity, particularly for species whose habitats and developmental success are tightly linked to thermal conditions (1–3).

Biological processes exhibit complex temperature dependencies across scales, from enzymatic catalysis to organismal growth. These dependencies are often represented by thermal performance curves, which display optimal performance at specific temperatures and declines at extremes. Such curves provide insights into how temperature governs life processes, ranging from enzymatic activity to embryonic development (4, 5). The Arrhenius equation (6–8) is widely used in biology to describe the temperature dependence of reaction rates. Initially derived for chemical reactions, it takes the form:

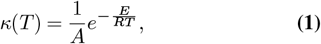

where *κ*(*T*) is the rate constant, *A* is a pre-exponential factor, *E* the activation energy, *R* the universal gas constant, and *T* the temperature in Kelvin. This response function is typically plotted as ln *κ*(*T*) versus 1*/T*, or using the associated time *τ* (*T*) = 1*/κ*(*T*):

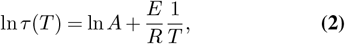

which yields a simple linear relationship. The Arrhenius equation predicts a monotonic increase in rate, which holds reasonably well over moderate temperature ranges (9, 10).

If the Arrhenius equation were universally valid and processes had similar activation energies, a few degrees of warming would speed up rates in a predictable way. For example, processes would speed up by ≈23% (assuming Q_10_ = 2, that is, the rate roughly doubles per 10^°^C). In reality, activation energies typically differ, so even under ideal Arrhenius kinetics different steps accelerate by differ-ent amounts and relative timings drift with temperature, potentially disrupting coordination ^1^.

Moreover, empirical observations show that even modest temperature increases often lead to pronounced deviations from Arrhenius behavior, especially at the upper end of the physiological temperature range. For instance, a temperature change of just a few degrees can trigger high fever in humans or completely arrest development in ectothermic embryos. In these non-Arrhenius regimes, biological rates frequently plateau, decline, or cease altogether. Such breakdowns arise from a range of factors, including protein denaturation, the activation of stress response pathways, and the nonlinear dynamics of complex biochemical networks (12, 16–18). These phenomena underscore the limitations of the Arrhenius framework in fully capturing biological temperature responses, particularly near physiological boundaries or across broad thermal ranges. They also highlight the need for mechanistic models that can explain not only species-level differences, but also variability across different realizations of biochemical networks.

To understand how such deviations appear throughout development, we examine temperature scaling across both early and later stages. Early embryonic cleavages are often driven by well-defined biochemical oscillators with minimal regulation, while later stages involve more complex and tightly regulated processes. Yet, as we show below, non-Arrhenius behavior occurs across this entire developmental spectrum, suggesting that it arises from general features of biological timing rather than stage-specific mechanisms.

Figures 1A and B illustrate this distinction. Panel A shows a schematic of the early, rapid cleavage divisions in the *X. laevis* frog embryo, regulated by a periodic cell cycle oscillator (12). These divisions occur without gap phases, checkpoint controls, or transcription (19, 20), allowing fast, regular cycles of proliferation. In contrast, Figure 1B sketches the slower and more variable timing of later developmental transitions, which involve diverse regulatory pathways, including gene expression, signaling, and morphogenesis.

**Fig. 1.**
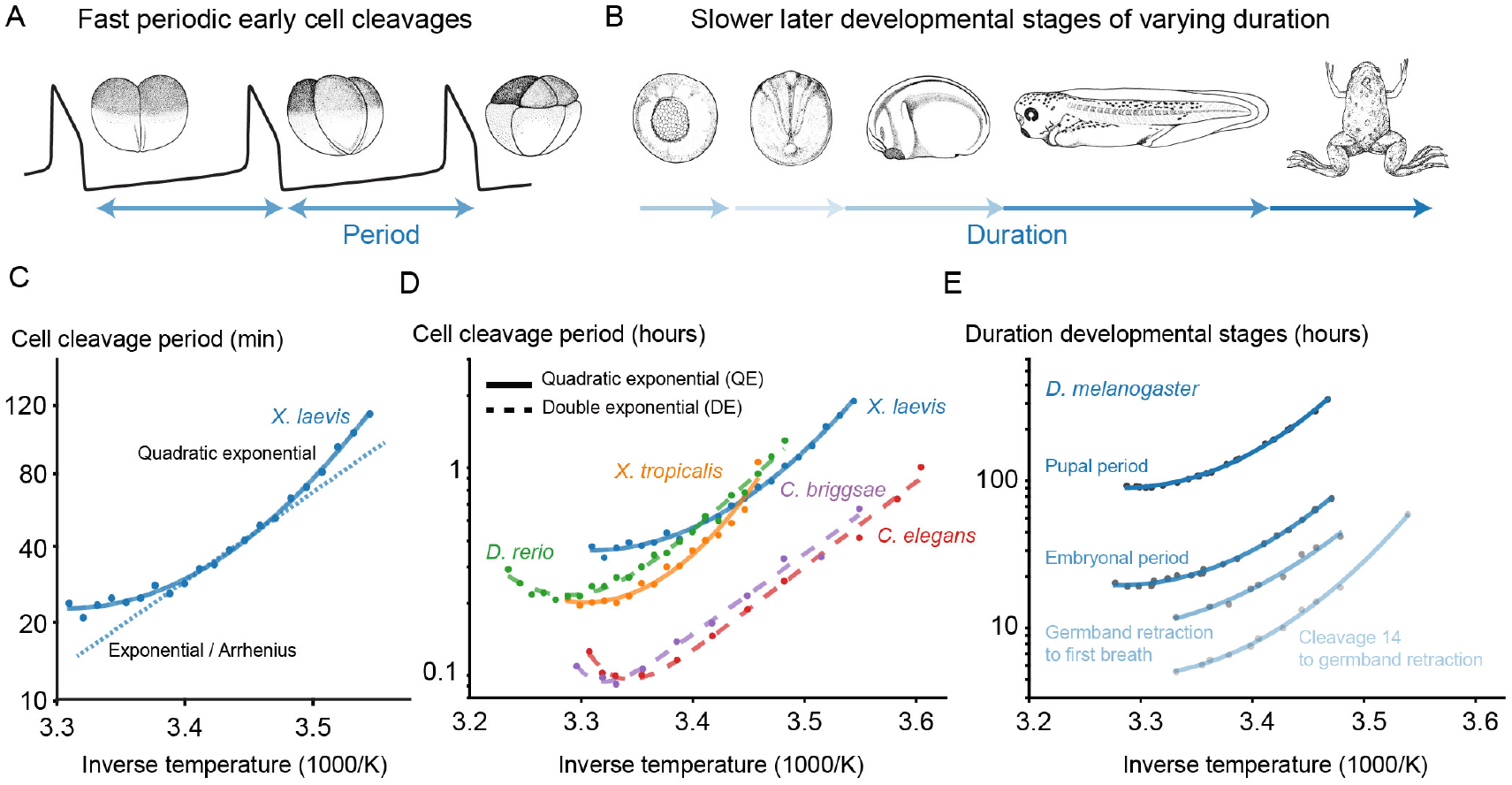
Developmental timing scales non-Arrhenius. (A) Schematic of the first cleavages in the early *X. laevis* embryo (adapted from (11)), driven by periodic cell cycle oscillations (12). (B) Schematic of selected later developmental stages during *X. laevis* development (adapted from (11)), showing varying and longer durations between stages. (C) Period of the early embryonic cell cycle in *X. laevis* (12), plotted on a logarithmic scale as a function of inverse absolute temperature. The best fits using the quadratic exponential (QE, solid line) and Arrhenius (dotted line) models are shown. (D) Periods of the early embryonic cell cycle for five species—*X. laevis, X. tropicalis, D. rerio* (12), and *C. briggsae, C. elegans* (13), with best fits using either the double exponential (DE, dashed line) or QE (solid line) models. (E) Timings of four developmental stages in *D. melanogaster*: from the 14th cell cleavage to germband retraction, germband retraction to first breath (14), total embryonic duration, and total pupal duration (15). Since both DE and QE models fit the data equally well, only the simpler QE fit (solid line) is shown.

Temperature scaling behavior across these developmental stages is illustrated in Figures 1C–E. In panel C, the early cell cycle period of *Xenopus laevis* is plotted on a logarithmic scale against inverse absolute temperature. The data appear approximately linear at lower temperatures, consistent with Arrhenius scaling, but deviate significantly at higher temper-atures. This upward curvature is more accurately described by the quadratic exponential (QE) model than by a single Arrhenius fit (Table 1). Panel D extends this analysis to five species: *X. laevis, X. tropicalis, D. rerio, C. briggsae*, and *C. elegans*, showing the temperature dependence of early cleavage timings. Most species display Arrhenius-like scaling at low temperatures, but deviate at higher temperatures. These deviations are abrupt in *C. elegans, C. briggsae*, and *D. rerio*, and more gradual in *Xenopus*. In some cases, the slope even becomes negative at high temperatures, implying negative local activation energies. The best-fitting model varies by species: the double exponential (DE) function fits the more abrupt deviations better, while the simpler QE law fits gradual ones equally well (Table 1). Panel E shows four later developmental intervals in *D. melanogaster*, including embryonic and pupal durations. These processes are more heterogeneous than early cleavages, yet they also deviate from Arrhenius scaling, especially over broader temperature ranges. Here too, the QE equation provides an excellent fit.

**Table 1.** Proposed modifications of the Arrhenius law for timings *τ*. Note that *E*_2_ can be negative.

| Scaling function | Form | Parameters |
| --- | --- | --- |
| Single-exponential (Arrhenius) | $Ae^{\frac{E}{RT}}$ | $A, E$ |
| Double-exponential (DE) | $A_1e^{\frac{E_1}{RT}} + A_2e^{\frac{E_2}{RT}}$ | $A_1, A_2, E_1, E_2$ |
| Quadratic-exponential (QE) | $Ae^{\left(\frac{E}{RT} + \frac{B}{T^2}\right)}$ | $A, B, E$ |

Together, these findings show that non-Arrhenius scaling is a general feature of developmental timing, appearing in both periodic and non-periodic processes, across diverse taxa and regulatory network architectures. The variability in scaling patterns between species and developmental stages highlights the need for a general mechanistic framework that can account for such differences.

To describe these deviations more accurately, several extensions to the Arrhenius model have been proposed. These include the QE equation (12, 14, 18, 21), the DE equation (13), the polynomial exponential (PE) equation (22), and others (23, 24). These models introduce additional temperature-dependent terms to improve data fits (see Supplementary Note 8). As shown in Figures 1C–E, both the QE and DE equations do a better job at describing cleavage and developmental timing than the Arrhenius equation. In species such as *C. elegans, C. briggsae*, and *D. rerio*, the DE law fits better due to abrupt deviations and negative local activation energies. In contrast, for *X. laevis, X. tropicalis*, and later stages of *D. melanogaster*, the QE law fits well with fewer parameters. These results suggest that different empirical models are suited to different biological contexts, depending on species, developmental stage, and underlying regulatory complexity.

Many of the models described above are empirical, aiming to fit the experimental data without necessarily explaining the underlying mechanisms. The Arrhenius equation itself was originally proposed as an empirical description of the exponential increase in reaction rates with temperature. Only later was a theoretical basis developed by Kramers (25, 26) and Eyring (27), who linked reaction rates to transitions over energy barriers in chemical reactions. More recently, we provided a mechanistic explanation for non-Arrhenius scaling in the early embryonic cell cycle of *Xenopus laevis*, using both embryos and egg extracts (12). By modeling the cell cycle as a relaxation oscillator, we showed that different activation energies among temperature-sensitive biochemical processes can account for the observed deviations from simple Arrhenius behavior. However, that framework is limited to oscillatory systems and does not extend to the temperature scaling of more complex, non-periodic biological processes. In complementary work, Voits and Schwarz (18) developed a statistical model that explains quadratic exponential scaling using a graph-based interpretation of mean first-passage times from a biochemical master equation. Such mechanistic approaches, drawing from dynamical systems and statistical physics, can help explain the physiological origins of temperature dependence in specific biological processes.

In this study, we present a general mechanistic framework based on Markov chains to explain non-Arrhenius temperature scaling in biological timing. We first map the ordinary differential equation (ODE) model for the cell cycle oscillator (12) to an equivalent Markov chain model, recovering the period as a mean first-passage time. We then show that temperature-dependent multi-step cascades produce a mean first-passage time with Arrhenius flanks and a central QE regime, the latter consistent with prior graph-theoretic work (18). Applying this framework to 121 published datasets spanning diverse rates, durations, and species, we consistently observe the Arrhenius–QE–Arrhenius pattern, with dataset-specific boundaries. The QE regime is a robust consequence of averaging across many steps, whereas the flanking Arrhenius regimes are governed by smaller subnetworks and vary more across realizations. Simple motifs (cycles, competing paths) also account for apparent negative activation energies. Together, the ODE-to-Markov bridge and cascade theory provide a unified, testable account of how network structure shapes thermal responses in development and related processes.

## Results

### The early embryonic cell cycle oscillator’s period is controlled by reaction-specific temperature sensitivities

#### A model based on coupled ordinary differential equations

The early embryonic cell cycle of the frog *Xenopus laevis* (and other fish and amphibians) functions as an autonomous oscillator, with oscillations driven by the interplay between Cyclin B synthesis, its regulated degradation, and the activation of Cyclin-dependent kinase 1 (Cdk1). These oscillations are governed by nonlinear interactions within a regulatory network, consisting of Cyclin synthesis (29), a bistable switch for Cdk1 activation (30, 31), and Cyclin degradation (32, 33) (Figure 2A). The simplicity of this system allows for mathematical modeling of the cell cycle using ODEs parameterized from experimental data (28, 34) (Supplementary note 1). These models successfully reproduce experimentally observed oscillations (Figure 2C), which form a closed trajectory, a limit cycle, in the (Cyc, Cdk1) phase plane (Figure 2D, red/blue). The limit cycle orbits around the system’s nullclines (brown), illustrating the dynamics of the cell cycle.

**Fig. 2.**
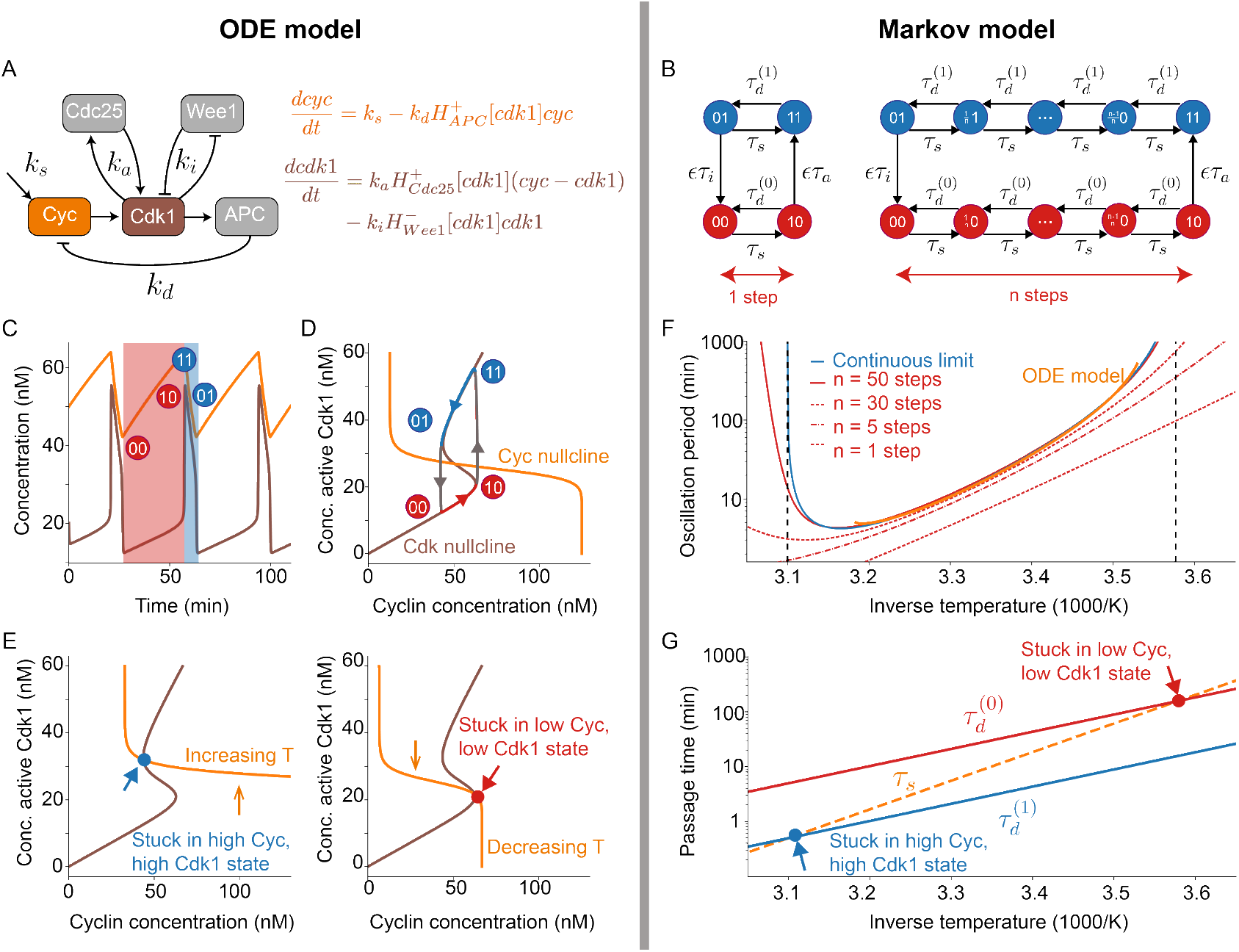
**The temperature scaling of** 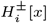 represent rising (+) and falling (−) Hill functions, which depend on the variable *x*. (B) Markov chain approximation of one cycle of the 2-ODE embryonic cell cycle model. The MFP time *E*[𝒯], calculated between the initial state (00) and the final state (00) (passing through all intermediate states in an anti-clockwise direction), is identified with the period of the cycle. (Left: one-step approximation) (Right: n-step approximation.) (C) A numerical simulation of the 2-ODE cell-cycle model, demonstrating a limit cycle as a time series, for a specific parameter set. (D) A numerical simulation of the 2-ODE cell-cycle model, demonstrating a limit cycle of the relaxation-oscillator type in phase space, for a specific parameter set. The nullclines are also drawn. (E) Illustration of how the shifting of the nullclines of the 2-ODE cell-cycle model with temperature can create new fixed points that cause the model to get stuck and destroy the limit cycle. (Left: increasing T) (Right: decreasing T). (F) Comparison of the period of the Markov model and the ODE model. Different step sizes and the continuous limit with “infinitely” small step sizes are shown. The ODE model, uses the same specific parameter set as the other figures. (G) Transition times 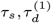 and 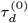 of the Markov model in function of temperature. The temperatures where the model gets stuck in a high Cyc and Cdk1 state, or in a low Cyc and Cdk1 state, correspond to asymptotes (dashed black lines) of the continuous limit in (F).

As the timescale of Cdk1 activation/inactivation is much faster than that of Cyclin synthesis and degradation, relaxation oscillations occur (35–41). During interphase, the system slowly ascends along the low-Cdk1-activity portion of the S-shaped nullcline, then abruptly jumps to the high-Cdk1-activity portion of the same nullcline (Figure 2D, red → blue curve). Subsequently, the system descends along the high-Cdk1-activity branch due to active APC/C-mediated Cyclin degradation, before falling abruptly back to the low-Cdk1-activity branch to complete the cycle (Figure 2D, blue →red curve). These dynamics produce characteristic sawtooth-shaped oscillations in Cyclin B levels (Figure 2C, brown).

This model has previously provided us with valuable information on the influence of temperature on the system (12). By introducing Arrhenius scaling to the rate constants (*k*_*s*_, *k*_*d*_, *k*_*a*_, and *k*_*i*_) (Supplementary note 1), the model allows to study how the oscillation period changes with temperature. When all activation energies (*E*_*a*_) are equal, the period exhibits Arrhenius-like scaling with a single *E*_*a*_. However, when activation energies differ across reactions, the model predicts deviations from Arrhenius scaling, particularly at high temperatures. These deviations, including negative effective activation energies, arise from the nonlinear interactions within the network and the differing temperature sensitivities of individual reactions.

Two distinct mechanisms were identified as drivers of non-Arrhenius scaling and thermal limits in embryonic cell cycles. First, thermal limits can occur when one or more reactions deviate from Arrhenius scaling, exhibiting a thermal optimum. In such cases, the system’s behavior is predominantly shaped by the dynamics of these biphasic reactions (12). Second, even when all reactions follow Arrhenius-like scaling, differences in activation energies among reactions lead to emergent thermal limits and deviations from linear Arrhenius plots. For example, if the activation energy for Cyclin synthesis is higher than that for Cyclin degradation, the ratio *k*_*s*_*/k*_*d*_ increases with increasing temperature. This causes the Cyclin nullcline to shift upward until it no longer intersects the middle portion of the S-shaped Cdk1 nullcline. At this point, oscillations cease, leaving the system in a M phase-like steady state with high Cdk1 activity (Figure 2E, left). Conversely, if temperatures decrease, the Cyclin nullcline shifts downward, stabilizing an interphase-like steady state with low Cdk1 activity (Figure 2E, right).

Strikingly, experiments using cycling *Xenopus* extracts revealed that both mechanisms contribute to overall temperature scaling. Cyclin synthesis exhibited a biphasic temperature response, and an imbalance in activation energies was also observed: Cyclin synthesis was more temperature-sensitive than degradation (12).

#### A model based on Markov jump processes

The ODE analysis shows that temperature scaling can be a network property. However, extending this analysis beyond relaxation oscillators remains a significant challenge. Boolean models motivate a discrete state space for regulatory networks (42), but their discrete time hinders analysis of temperature-dependent periods. We therefore map the two-ODE oscillator to a continuous-time Markov jump process on a discrete state graph with protein concentrations having binary states: active (1) or inactive (0). We introduce a continuous-time framework into the state graph by assigning transition rates *κ*_*ij*_ to transitions between states (*i*) → (*j*). For small time intervals *dt*, the probability of such a transition is given by:

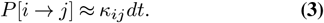

This describes a Markov jump process, a stochastic, continuous-time process on a discrete state space with minimal memory (43). The future state of the system depends only on its current state, making these processes both analytically and numerically tractable.

In a Markov process, the time required to transition from state *i* to state *j* is called the first passage time 𝒯 _*ij*_. We are especially interested in the first passage time from the initial state 1 to the final state *n*, denoted as 𝒯 _1*n*_ = 𝒯. As 𝒯 varies between realizations, it is treated as a random variable. However, the mean first passage (MFP) time 𝔼 [𝒯] a fixed value for the process, can be calculated and used to study biological transition timings. For simplicity, we assume that the higher order moments of 𝒯 are negligible compared to the mean, a reasonable approximation for large-scale biological systems evolved for specific purposes.

Temperature dependence is introduced into the Markov jump process by assigning Arrhenius scaling to the transition rates. Specifically, the inverse rate 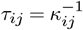 is modeled as:

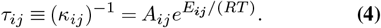

where *A*_*ij*_ is the pre-exponential factor, *E*_*ij*_ is the activation energy, *R* is the universal gas constant, and *T* is the absolute temperature in Kelvin.

To approximate the two-ODE model, we developed a simplified Markov chain in which Cyclin B and active Cdk1 are limited to binary states: ON (1) or OFF (0) (Supplementary note 3). The ON state represents a high concentration of Cyclins and high Cdk1 activity, while the OFF state corresponds to a low concentration of Cyclins and low Cdk1 activity. This yields four possible system states: (0, 0), (1, 0), (1, 1), and (0, 1). Transition times *τ*_*ij*_ were assigned based on the corresponding rate constants in the two-ODE model. For example, *τ*_*s*_ represents the time for Cyclins to be synthesized [transitions (0, 0) → (1, 0) and (0, 1) → (1, 1)], while *τ*_*d*_ governs Cyclin degradation [transitions (1, 0) → (0, 0) and (1, 1) → (0, 1)]. Similarly, *ϵτ*_*a*_ and *ϵτ*_*i*_ were used for the faster activation and inhibition of Cdk1, leading to transitions (1, 0) → (1, 1) and (0, 1) → (0, 0), respectively. To replicate the two-rate dynamics of Cyclin degradation, we introduced two pre-exponential factors, 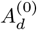 (low degradation when APC is OFF) and 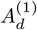 (high degradation when APC is ON). These factors were scaled to match the ratio of extremal values in the Hill function of the two-ODE model.

The resulting Markov process is represented as a weighted directed graph (Figure 2B, left). While backward reactions are possible, they were omitted in the transitions for Cdk1 activation and inhibition for simplicity. At a given temperature, the MFP time for the system to traverse all states in an counterclockwise direction, starting from (0, 0) and returning to (0, 0), was calculated and identified as the period of the cycle. The results were then compared to the temperature-dependent period of the two-ODE model. To improve the match between the Markov chain and two-ODE models, intermediate states were added for Cyclin B. Instead of binary states, Cyclin B concentrations were divided into *n* states, allowing transitions in steps of 1*/n*. The right-hand side of Figure 2B shows the resulting state graph. As *n* increases, the MFP time of the Markov chain converges to the period of the

ODE model (Figure 2F). In the continuous limit (*n*→∞), the system exhibits vertical asymptotes at extreme temperatures (dashed lines), corresponding to thermal limits where oscillations cease.

The Markov model provides an intuitive explanation for the breakdown of Arrhenius scaling when *E*_*s*_ ≠ *E*_*d*_. Because transitions are stochastic in both directions, net forward drift is observed when the mean waiting times satisfy:

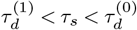

When *E*_*d*_ and *E*_*s*_ differ, their scaling with temperature diverges, causing one of these inequalities to fail at extreme temperatures (see Figure 2G). This results in the system becoming trapped in backward cycles during either Cyclin synthesis or degradation, halting oscillations (when *n* → ∞). Physically, this corresponds to one process dominating the other, preventing the system from completing a full cell cycle. In short, the Markov formulation reproduces the ODE period and the loss of oscillations at thermal extremes, while making a forward–backward bias explicit. This, in turn, motivates the three-regime scaling we derive next for linear cascades.

#### Cascades of reversible transitions explain temperature scaling in developmental systems

Motivated by the ODE to Markov mechanism above, we now examine linear cascades of reversible steps as a simplified model of developmental progression. This shifts the focus from periodic cycles to one-way progressions, where a system transitions from an initial to a final state through a series of potentially reversible steps. Many later developmental events can be conceptualized this way, as sequential, partly reversible processes rather than a dedicated oscillatory mechanism.

To generalize this framework, we allow the transition times between states to differ. The resulting state graph, a linear cascade of cycles, is shown schematically in Figure 3A. It consists of *n* states, where each state *i* connects to the next state *i* + 1 with a forward transition time *τ*_*i,f*_ and to the previous state *i* − 1 with a backward transition time *τ*_*i,b*_. ^2^ The process begins in state 1 and terminates in state *n*. For this linear cascade, the MFP time from 1 to *n* over all realizations of the Markov process can be calculated analytically:

**Fig. 3.**
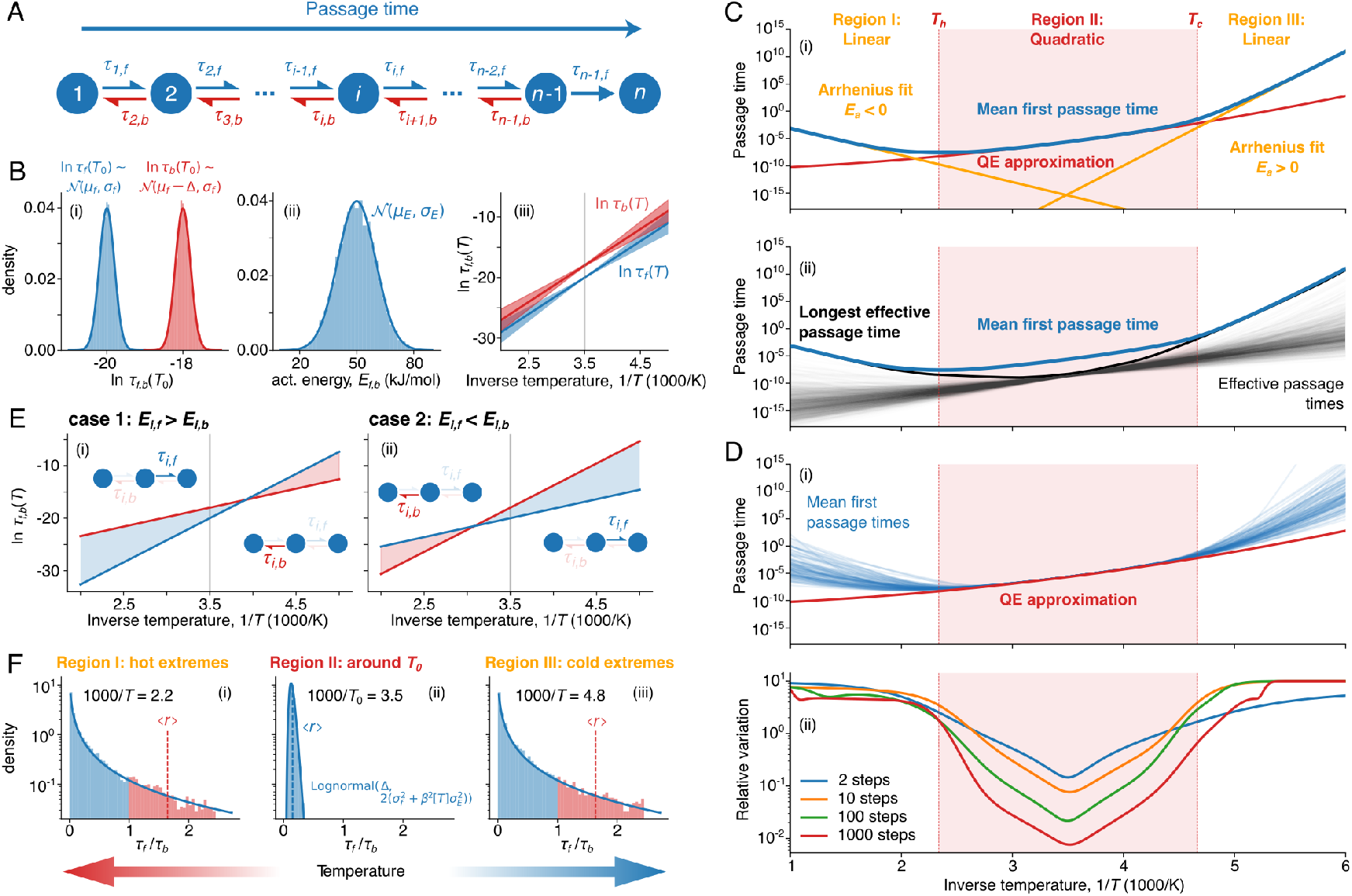
The mean first passage time of a linear cascade of reversible processes exhibits three distinct temperature-scaling regimes. (A) Schematic of a linear cascade of reversible Markov jump processes. (B) Ingredients of the transitions times’ temperature dependence: (i) natural logarithms of forward (blue) and backward (red) transition times are distributed normally at the reference temperature *T*_0_, with the backward distribution shifted by Δ with respect to the forward one; (ii) activation energies of forward and backward transitions are drawn from normal distribution with the same mean and variance; (iii) resulting Arrhenius scaling of forward (blue) and backward (red) transition times (mean±s.d.). Gray vertival line indicates inversed reference temperature (1*/T*_0_). (C) Simulation of the temperature dependence of the MPF time for system (A): (i) Arrhenius scaling is fitted at high and low temperatures (Regions I and III, yellow lines). In Region II, the QE approximation from Eq. (7) provides a good fit. Boundaries of the QE region are determined from Eq. (8). (ii) MPF time plotted alongside with effective passage times (family of thin black curves) and the longest effective passage time (bold black curve). (D) Variation of MPF time temperature scaling across multiple random realizations of system (A): (i) MPF time from 100 realizations are plotted alongside with the QE approximation; (ii) Relative variation of MPF time (s.d.*/*mean) across realizations versus temperature for different sizes of a linear cascade (A). (E) Examples of individual transition time scaling when *E*_*i,f*_ *> E*_*i,b*_ (i) and *E*_*i,f*_ *< E*_*i,b*_ (ii). In both panels, blue and red lines show scaling of individual forward and backward transition line, respectively; blue and red shadings show the areas where either forward or backward transition from state *i* is more preferable, as shown by the respective pictograms. (F) Forward-to-backward transition time distribution illustrating the fraction of forward-dominated (*r <* 1, blue bars) and backward-dominated (*r >* 1, red bars) transitions, at different temperatures symmetric to the reference one.

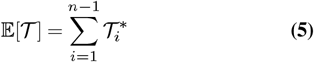

where

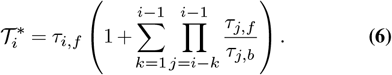

When forward transitions dominate (i.e., *r*_*i*_ = *τ*_*i,f*_ */τ*_*i,b*_ ≪1), the MFP time simplifies to the sum of all forward transition times, 𝔼 [𝒯] ≈ *τ*_*i,f*_. However, when backward transitions become significant *τ*_*i,b*_ = 𝒪 (*τ*_*i,f*_) or *τ*_*i,b*_ *< τ*_*i,f*_, 𝒯 increases, as each effective passage time 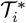 is amplified by a factor that accounts for the ratios of forward and backward transitions in prior cycles. This factor is always greater than 1, meaning faster backward transitions increase the MFP time considerably.

To model large-scale biological processes, we fix a reference temperature *T*_0_ and generate transition statistics in three steps (see also Supplementary Note 5).

i. *Baseline times at T*_0_: for each step *i*, we draw for-ward transition rates ln *τ*_*i,f*_ (*T*_0_) from 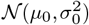 and back-ward transition rates ln *τ*_*i,b*_ (*T*_0_) from 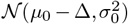, where 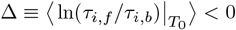 sets a forward bias at *T*_0_ (forward faster than backward), see Fig. 3B(i).
ii. *Activation energies:* independently for each directed transition, we draw activation energies *E*_*i,f*_ and *E*_*i,b*_ from 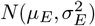, with biologically plausible values (e.g., *µ*_*E*_ ≃ 50 kJ mol^−1^, *σ*_*E*_ ≃ 10 kJ mol^−1^; Ref. (44); see Fig. 3B(ii)).
iii. *Arrhenius extrapolation from T*_0_: for any temperature *T*, the transition times are given by

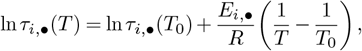

which yields the forward/backward (= *f/b*) time distributions shown in Fig. 3B(iii).

In the limit of many steps (*n* → ∞) this leads to a QE scaling law (Eq. (70) of the Supplementary note 5):

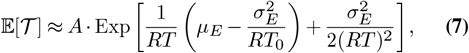

where *A* is a prefactor determined by the baseline statistics at *T*_0_ (and depends on *n, µ*_*E*_, and *σ*_*E*_). Although illustrated here with normally distributed activation energies, the QE form is generic, arising for broad classes of energy distributions beyond normality. The QE scaling law is valid within a specific temperature range:

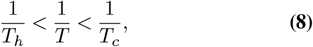

where forward transitions dominate, meaning ⟨*r*⟩ = ⟨*τ*_*i,f*_ */τ*_*i,b*_⟩ < 1. Here, *T*_*c*_ and *T*_*h*_ are the temperature limits of the QE scaling at cold and warm temperatures:

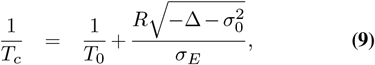

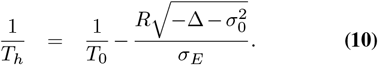

Note that the above expressions for the temperature limits *T*_*h*_ and *T*_*c*_ only have a physical meaning for 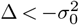, i.e., when the distributions of forward and backward transition times at the reference temperature are well-separated.

Within this range, the MFP time follows a QE scaling law as described in Eq. (7) (Figure 3C). This behavior arises from the collective contribution of many forward-dominated transitions with comparable timescales in the sum given by Eq. (5). As a result, the system displays strongly reduced variability across realizations, particularly as the number of steps increases (Figure 3D, Figure S3). This regime is therefore both more predictable and more robust, reflecting the emergent regularity of large ensembles of similarly activated steps. The validity of the QE regime largely depends on the relative width of the activation energy distribution and mean directional bias |Δ|.

In contrast, outside this range, the distribution of effective transition times 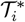 includes large tails, outliers dominate the total MFP time, and the law of large numbers is no longer valid (Figure 3C). In these extreme temperature regions, the MFP time scales in an Arrhenius-like manner with either a positive or negative constant activation energy, depending on the temperature limit: positive activation energy in the cold regime and a negative activation energy in the warm regime.

In both cases, the dynamics are effectively dominated by the longest rate-limiting effective passage time 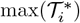 (panel (ii) in Figure 3C). This leads to strong sensitivity to stochastic variation: despite identical distributions of activation energies across realizations, substantial variability emerges in the observed slopes, transition sharpness, and crossover temperatures. This variability persists even for large systems (large *n*), indicating that the statistical averaging in long cascades does not apply in these regimes (Figure 3D, Figure S3).

To better understand where the deviations from the QE scaling law originate, let us zoom into the scaling of individual reactions. Since the activation energies of forward and backward transitions of individual reaction are generally unequal, two cases are possible: (1) *E*_*i,f*_ *> E*_*i,b*_ and (2) *E*_*i,f*_ *< E*_*i,b*_. In both cases, forward transitions are preferable around the reference temperature *T*_0_, favoring the QE approximation.

However, backward transitions in case (1) dominate at lower temperatures, while the opposite holds for the case (2) (Figure 3C, Figure S2, and Supplementary note 5). Thus, moving away from *T*_0_ in either direction facilitates the backward-dominated transitions with *r*_*i*_ = *τ*_*i,f*_ */τ*_*i,b*_ *>* 1 through either case-(1) reactions on the colder side or case-(2) reactions on the warmer side (Figure 3F). Such backward-dominated transitions trap the cascade in the local cycle motifs, thereby extending overall MFP time beyond the QE prediction at the temperature extremes.

Together, these results show that complex biological processes can transition between distinct scaling regimes (Figure 4A). Quadratic exponential scaling arises in intermediate thermal regimes due to cumulative contribution of many steps in large cascades, while single-exponential (Arrhenius) behavior dominates at thermal extremes due to the disproportionate influence of a small number of transitions. This framework provides a mechanistic basis for understanding both the robustness and variability observed in temperature-dependent biological timing.

**Fig. 4.**
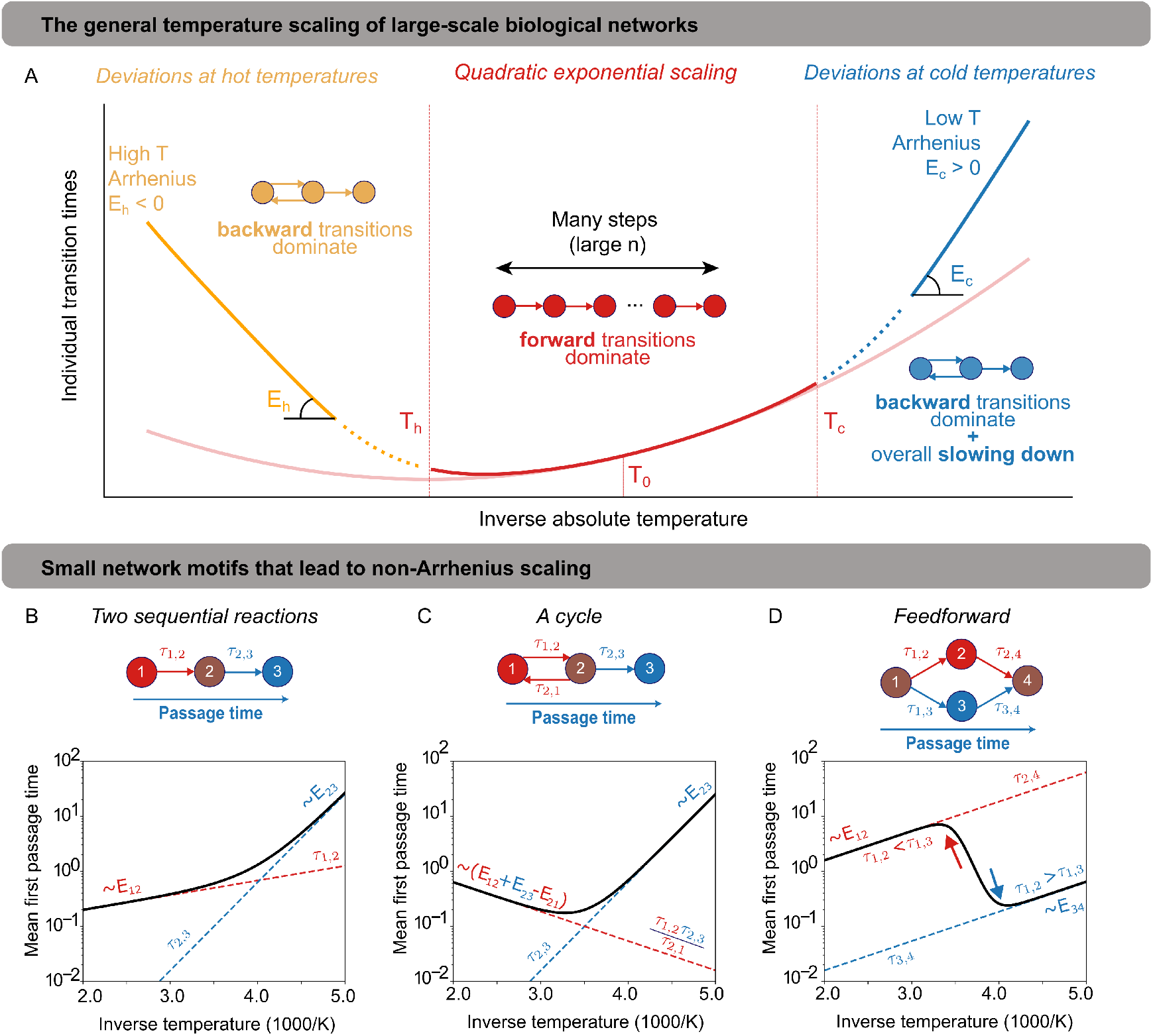
General scaling motifs underlying temperature-dependent biological timing. (A) Schematic overview showing how biological systems combine quadratic exponential scaling at intermediate temperatures — driven by the collective averaging of many similar transitions — with Arrhenius-like single-exponential scaling at low and high temperature extremes, where one or few rate-limiting steps dominate. (B) Sequential two-step motif: at low temperatures, the slower high-barrier reaction sets the overall rate; at high temperatures, the faster low-barrier reaction dominates, but both regimes maintain positive effective activation energies. (C) Cycle motif: at high temperatures, reverse transitions become dominant, trapping the system and producing an emergent negative macroscopic activation energy despite all microscopic steps following Arrhenius scaling. Here, *E*_23_ *> E*_12_ and *E*_12_ + *E*_23_ *< E*_21_. (D) Feedforward motif: two parallel forward pathways contribute, with temperature shifting dominance between them, producing a negative activation energy within a narrow intermediate temperature range.

### Simple network motifs explain the mechanisms behind non-Arrhenius scaling and negative activation energies

Figure 4A illustrates how single-exponential Arrhenius behavior, whether with positive or negative activation energies, dominates at thermal extremes, due to the disproportionate influence of a small number of rate-limiting transitions. We focus here on three minimal network motifs that play a critical role in generating deviations from simple Arrhenius responses at these temperature extremes.

The simplest case is two sequential reactions with different activation energies (Figure 4B). At low temperatures, the system’s overall rate scales with the reaction having the largest activation energy, while at high temperatures, it scales with the smallest. This produces the widely used double-exponential response curve. However, at both extremes, the system still exhibits scaling with a positive activation energy. So how can minimal network motifs, where each microscopic transition obeys the Arrhenius law with a fixed positive activation energy, give rise to a system where the MFP time scales with a negative activation energy?

We identify two minimal state-graph motifs that illustrate how negative activation energies can emerge from systems with only positive microscopic activation energies (see Supplementary Note 7). The first is a simple cycle (Figure 4C), where the system can transiently loop between states 1 and 2 before reaching the endpoint 3. At high temperatures, when reverse transitions become significantly faster than forward ones, the MFP time scales with an effective negative activa-tion energy. The second motif involves two parallel forward-directed paths (Figure 4D), where temperature determines the likelihood of taking one path over the other. Within a narrow intermediate temperature range, the shift in path dominance produces a negative activation energy.

These motifs demonstrate how macroscopic negative activation energies can arise from underlying microscopic processes that individually follow Arrhenius scaling, emphasizing the importance of network topology and transition probabilities in shaping overall temperature dependence. Crucially, both motifs feature multiple possible routes to the endpoint, with the most probable path shifting with temperature. In contrast, systems with only a single possible path, such as the two-step sequential motif, cannot generate negative macroscopic activation energies in their MFP scaling.

The temperature-dependent scaling in the cycle and feed-forward motifs is qualitatively distinct. For the cycle, the macroscopic activation energy can become negative in the high-temperature limit, as the system becomes increasingly trapped in reverse loops. For the feedforward motif, negative activation energies emerge only in an intermediate transition region, where the dominant contribution to the MFP time shifts between two competing forward paths that control behavior at opposite thermal extremes.

#### The Markov framework captures temperature scaling across diverse species, biological timescales, and temperature ranges

To evaluate the applicability of the Markov cascade model, we assembled a diverse set of experimental datasets densely sampled in temperature, i.e. many measurements per curve across a broad thermal span, covering growth and developmental processes, including early embryonic cell cycles and stage transitions across multiple species (12–15, 45–88). In total, we analyzed 121 datasets from 48 sources, comprising 2174 measurements with ≈18 temperature points per curve (and minimum 5), with a typical temperature span of 30 ± 17 K (minimum 10K).

Each temperature response curve was fit with a composite model: a central QE regime [Eq.(7)], flanked by single-exponential regimes at low and high temperatures (See Supplementary note 8). This approach enables a unified description of thermal scaling across biologically relevant ranges, with transitions between regimes governed by the underlying distribution of activation energies and the balance of forward and backward transitions in the Markov chain. Figure 5A shows representative examples, highlighting the diversity of observed scaling behaviors, from processes following QE across the full range to cases with deviations at either thermal extreme. Across all datasets, the model provided excellent fits (median *R*^2^ = 0.98, 90% CI [0.89, 0.99]), accurately capturing both gradual and sharp transitions in scaling. The majority of experimental temperature responses, approximately 59%, were fully described by the QE law, while the rest displayed deviations at temperature extremes: 32% only at warm temperatures, 6% only at cold temperatures, and 3% at both ends (Figure 5B). This pattern suggests that while many biological systems operate within a robust QE regime near their thermal midpoint, transitions to Arrhenius-like scaling are common at extreme temperatures. At the same time, datasets exhibiting deviations at colder temperatures were less represented among the analyzed sources. It may reflect the fact that the processes slow down faster at the colder side, making their experimental measurement more challenging.

**Fig. 5.**
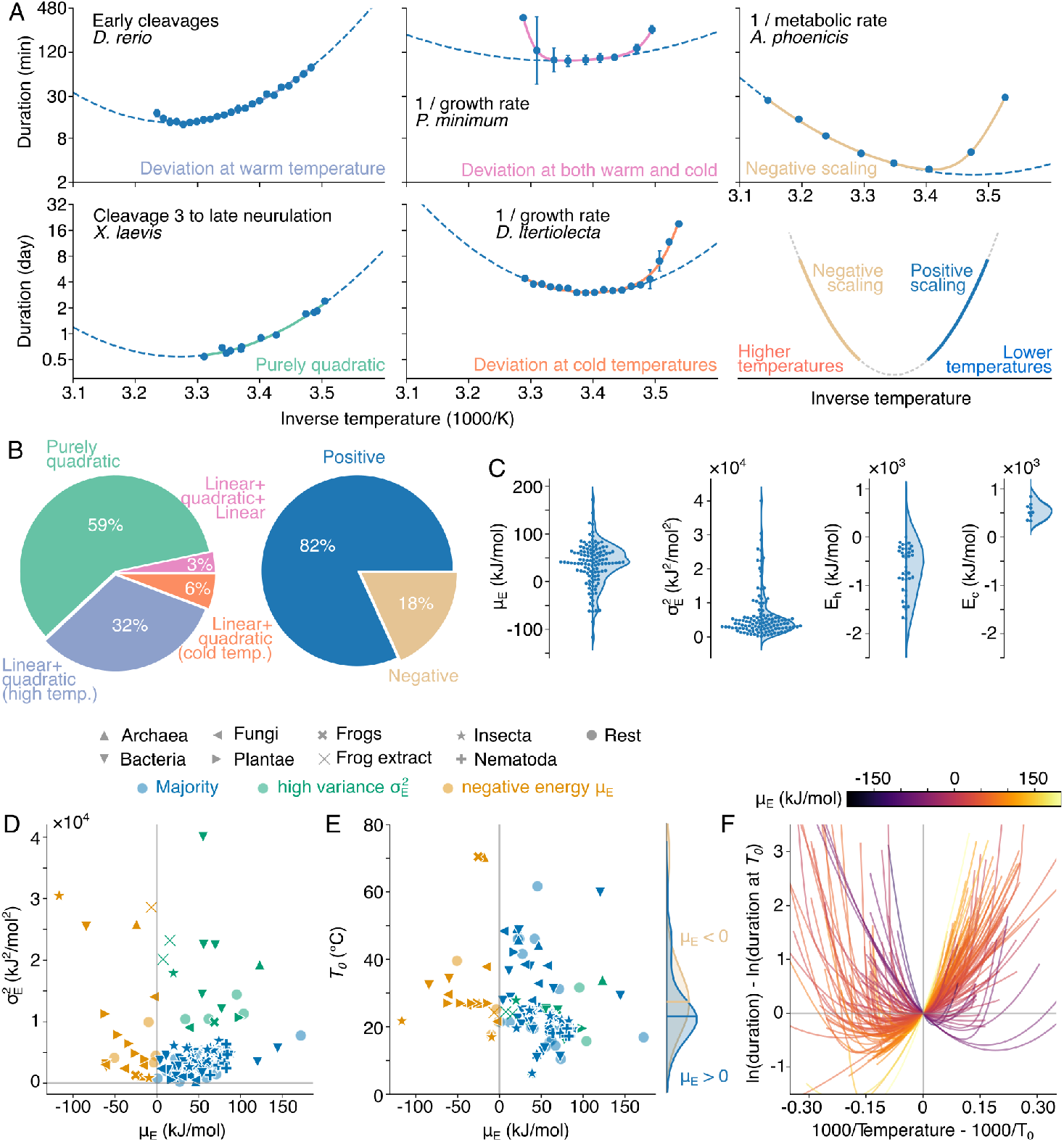
The Markov cascade model explains temperature scaling across diverse biological systems. (A) Example fits of experimental data using a composite model with a central quadratic exponential (QE) regime flanked by exponential regimes at low and high temperatures. (B) Proportions of *N* = 121 datasets classified as fully QE or showing deviations at warm, cold, or both extremes (left chart) and as positive or negative scaling (right chart). (C) Distributions of fitted QE parameters, including mean activation energy *µ*_*E*_, variance 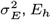 and *E*_*c*_. (D) Scatterplot of datasets in 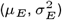 space, with markers indicating taxonomic groups (see Legend) and colors indicating scaling regime (QE or deviations, similar as in B). (E) Scatterplot similar as in (D) but in (*µ*_*E*_, *T*_0_) space. Kernel density estimation compares reference temperature *T*_0_ in datasets with positive and negative scaling (*p <*.01 via the Mann-Whitney test; solid lines indicate sample medians). (F) Temperature response curves normalized to their value at reference temperature *T*_0_, overlaid and colored by *µ*_*E*_, showing conserved yet distinct scaling behaviors across datasets.

Figure 5C presents the distribution of key fitted parameters. The mean activation energy of the QE regime, *µ*_*E*_ (Eq. 7), typically ranged from 40–80 kJ/mol, consistent with systems composed of many elementary steps, each drawing from a moderate activation energy distribution (0–200 kJ/mol). These observed mean activation energy values also overlap those reported by Dell *et al*. (10). The width of the distri-bution, 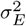, varied across systems and determined the size of the QE window as well as the scaling curvature. For temperatures outside this window, we extracted effective activation energies from the flanking exponential regimes, confirming that distinct scaling laws govern different parts of the thermal response.

We further visualized these relationships in the 2D parame-ter space 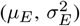, shown in Figure 5D. Here, we marked different major clades (Archaea, Bacteria, Fungi, and Plan-tae) with different symbols, and indicate data from frogs (including frog egg extracts), Insecta, Nematoda, and all other sampled groups separately. Most datasets cluster around a positive mean activation energy *µ*_*E*_ with moderate variance 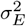. Two anomalous groups emerge: (i) systems with exceptionally large variance 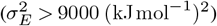 and (ii) systems with a *negative* mean activation energy of the QE regime (*µ*_*E*_ *<* 0). These outliers are enriched for phytoplankton, algae, bacteria, viruses, thermophilic fungi, and heat-tolerant plants, taxa that routinely experience severe thermal conditions. This pattern suggests distinct temperature-compensation strategies: a large 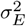 is consistent with stronger compensation across the operative thermal window, whereas the biological interpretation of *µ*_*E*_ *<* 0 is less direct.

The plot of the reference temperature *T*_0_ against *µ*_*E*_ (Figure 5E) shows that the *µ*_*E*_ *<* 0 group has a higher median *T*_0_ by ~4.5 ^°^C relative to the population with *µ*_*E*_ *>* 0, consistent with a change of the operating range towards warmer conditions. Consistently, responses normalized by *T*_0_ and colorcoded by *µ*_*E*_ (Figure 5F) reveal that *µ*_*E*_ sets the local slope at *T*_0_, effectively sliding systems along the QE curve: positive *µ*_*E*_ aligns with the cooler-temperature side, whereas *µ*_*E*_ *<* 0 aligns with the warmer-temperature side (see schematic in Figure 5A).

#### Negative quadratic exponential temperature scaling can be recapitulated in the Markov cascades

Data reveal two distinct settings in which the apparent activation energy can be negative: (i) at the *hot extreme*, where reverse loops in cyclic motifs dominate and the macroscopic slope becomes negative; and (ii) within the *intermediate QE range*, where a local negative slope is observed over a finite window. The hot-edge case has a simple mechanistic account via reverse-loop trapping in cycles (cf. Fig. 4C). In contrast, the mechanism in the QE window case is unclear. We therefore explore thermodynamically consistent network options, where all microscopic steps obey Arrhenius with positive activation energies, showing how negative apparent *E*_*a*_ can appear within the QE window.

The most direct way to reproduce negative scaling is to implement a linear Markov cascade in which the activation energy distribution is mirrored around zero (Figure 6A–C). Drawing microscopic one-step transition energies from this distribution generates negative linear scaling of individual reactions, producing a negatively scaled temperature response curve that qualitatively resembles those observed in the experimental datasets (Figure 5A, F). However, interpreting a negative mean activation energy simply as the average over negative microscopic transition energies can be misleading, since such a perspective may contradict fundamental thermodynamic principles.

**Fig. 6.**
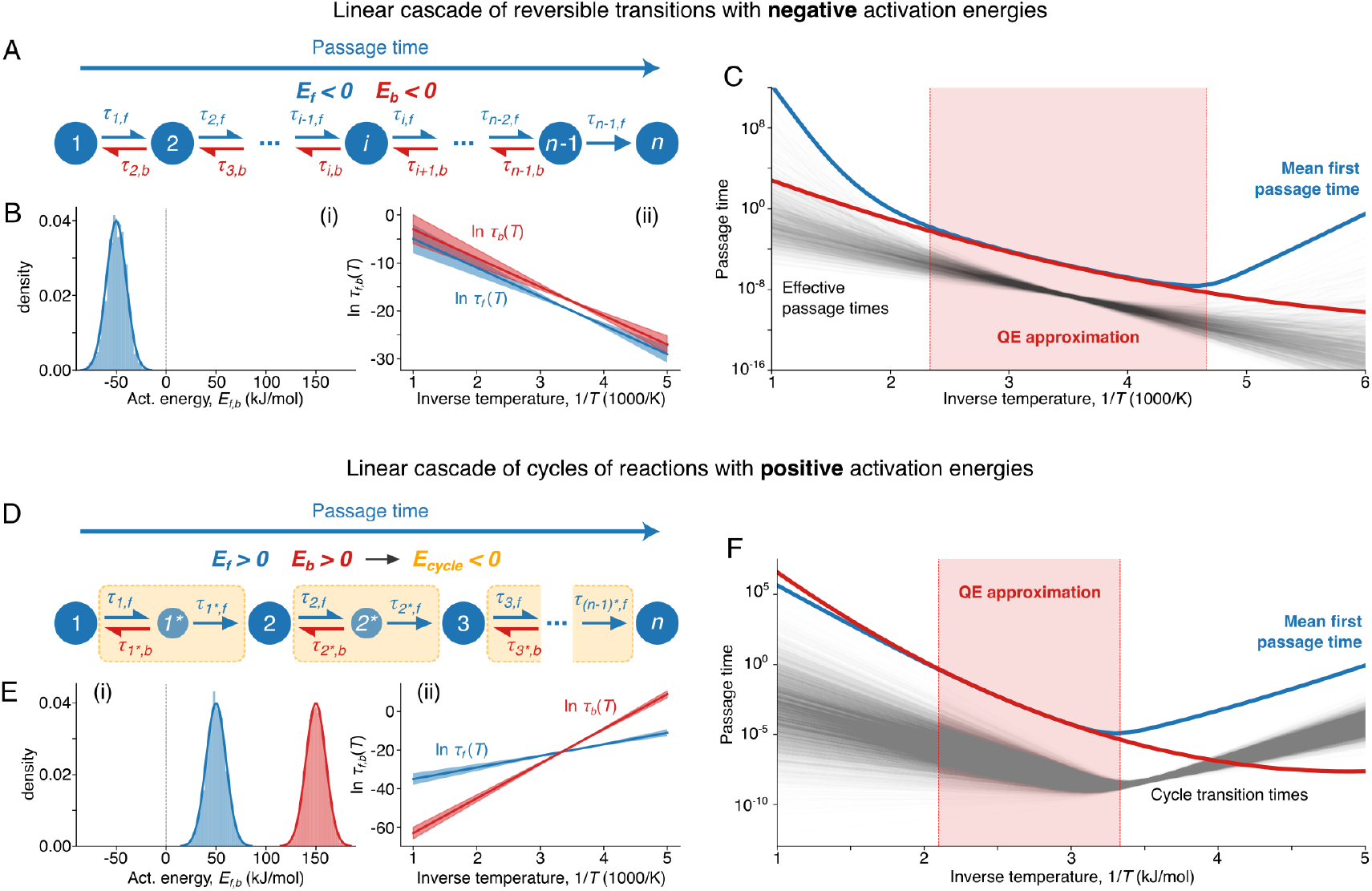
Negative quadratic-exponential temperature scaling can be recapitulated in the Markov cascade of cycles. (A)-(C) Negative MFP time temperature scaling results from the negative scaling of individual reactions in a linear Markov cascade. (A) Schematic of a linear cascade, where each reaction has negative activation energy (*E*_{*f,b*}_ *<* 0). (B) Temperature scaling of individual reactions: (i) normal distribution from which activation energies *E*_{*f,b*}_ are drawn; (ii) “negative” scaling of individual forward and backward reactions in blue and red, respectively (mean±s.d.). (C) Resulting temperature scaling of MFP time in the linear cascade (blue). The bold red curve shows QE approximation, the thin black curves display effective passage times, and the shading indicates the range where MFP time scaling follows the QE approximation. (D)-(F) Cascade of cycles with an apparent negative scaling branch composed of positively scaled reactions can also exhibit negative MFP time scaling. (D) Schematic of a linear cascade of cycles, where each reaction has positive activation energy (*E*_{*f,b*}_ *>* 0), yielding negative apparent activation energy of cycles (*E*_*cycle*_ *<* 0). (E) Temperature scaling of individual reactions: (i) normal distributions from which activation energies *E*_{*f,b*}_ are drawn; (ii) “positive” scaling of individual forward and backward reactions in blue and red, respectively (mean±s.d.). (F) Resulting temperature scaling of MFP time in the linear cascade of cycles (blue). The bold red curve shows QE approximation, the thin black curves display individual cycle transition times, and the shading indicates the range where MFP time scaling follows the QE approximation.

We propose instead that systems with negative *µ*_*E*_ may reflect a higher degree of regulation, in which transitions between states are multi-step and can thus give rise to negative *apparent* activation energies. To illustrate this, we turn to minimal network motifs composed solely of positive-energy transitions, for which we have previously demonstrated the possibility of negative scaling (Figure 4B–D). Without loss of generality, we focus on a cycle motif and consider a Markov cascade of such cycles (Figure 6D). In this setup, forward and backward one-step transitions, each positively scaled but dif-fering in mean activation energies (Figure 6E), together produce negative scaling of individual cycles at higher temperatures and positive scaling at lower temperatures (Figure 6F). As expected, the MFP time of a cascade of such cycles then displays negative QE scaling within a defined temperature range under physically realistic conditions. The simple cycle examined here serves merely as one example, as many other multi-step motifs could plausibly underlie negatively scaled reactions.

## Conclusion

Temperature fundamentally shapes biological timing. While the Arrhenius equation captures how individual activated steps accelerate with temperature, its limits become clear across broader thermal ranges and complex multi-step processes. Here, we introduced a Markov cascade framework that provides a mechanistic account of these departures.

Across linear cascades of reversible steps, the MFP time exhibits a consistent three-zone pattern: a QE regime at intermediate temperatures, with Arrhenius-like behavior at cold and hot extremes. The QE regime arises from statistical averaging over many temperature-sensitive steps and is robust to distributional details, yielding predictable and low-variability timing. At thermal extremes, averaging breaks down and a few effective steps dominate, restoring single-exponential (Arrhenius) scaling and increasing variability. We derived analytical boundaries between these regimes in terms of activation energy statistics and forward-backward bias, and we used an ODE → Markov mapping of the embryonic cell-cycle oscillator to reproduce the observed thermal limits of oscillations.

Apparent negative activation energies need not imply negative microscopic barriers or enzyme denaturation. In our framework they emerge naturally from network topology. Simple motifs, such as cycles or feedforward parallel pathways, are sufficient to generate such effects, emphasizing how global thermal responses can emerge from local, reversible steps.

The theory agrees with data: applied to more than 100 densely sampled datasets spanning taxa, rates, and developmental intervals, the composite QE–Arrhenius picture provides excellent fits, explains when coordination is preserved, and clarifies why breakdowns can occur near physiological limits. These results offer practical guidance for measurement (need for dense temperature sampling) and interpretation (treat warm-edge slowdowns and negative slopes as network effects unless denaturation is independently demonstrated), and they suggest how thermal robustness may be organized by network architecture in natural systems.

Our QE prediction is consistent with graph-theoretic first-passage analyses of large networks (18). We specify when QE holds for *finite* linear reversible cascades, explain the reemergence of Arrhenius scaling at the extremes and motif-driven negative apparent activation energies, and validate these predictions across a broad empirical dataset.

More broadly, the Markov cascade model provides a unified, testable framework for forecasting how biological timing shifts with temperature. By linking microscopic activation statistics to macroscopic timescales, it identifies thresholds of developmental failure, guides targeted perturbations (e.g., altering network topology or biases), and can be extended beyond development to other multi-step physiological and ecological processes in a warming world.

## Discussion

### Robustness of biological timing to temperature changes

#### Development

Embryonic development is a cascade of coordinated biochemical events, from molecular regulation to large-scale morphogenesis. Despite large changes in stage duration with temperature, the final outcome often remains remarkably consistent. A striking example is *Drosophila*, where the relative durations of embryonic and pupal stages are preserved across a broad thermal range (14, 70) (Figure 7A). Similar invariance is seen in *X. laevis* embryos (12, 14) (Figure 7B) and in Medaka fish somitogenesis, where somite spacing remains constant even though the period varies more than threefold (89). Related results show that transcriptional output and cell cycle timing scale proportionally in early *Drosophila* embryos (90), preserving gene expression levels across temperatures.

**Fig. 7.**
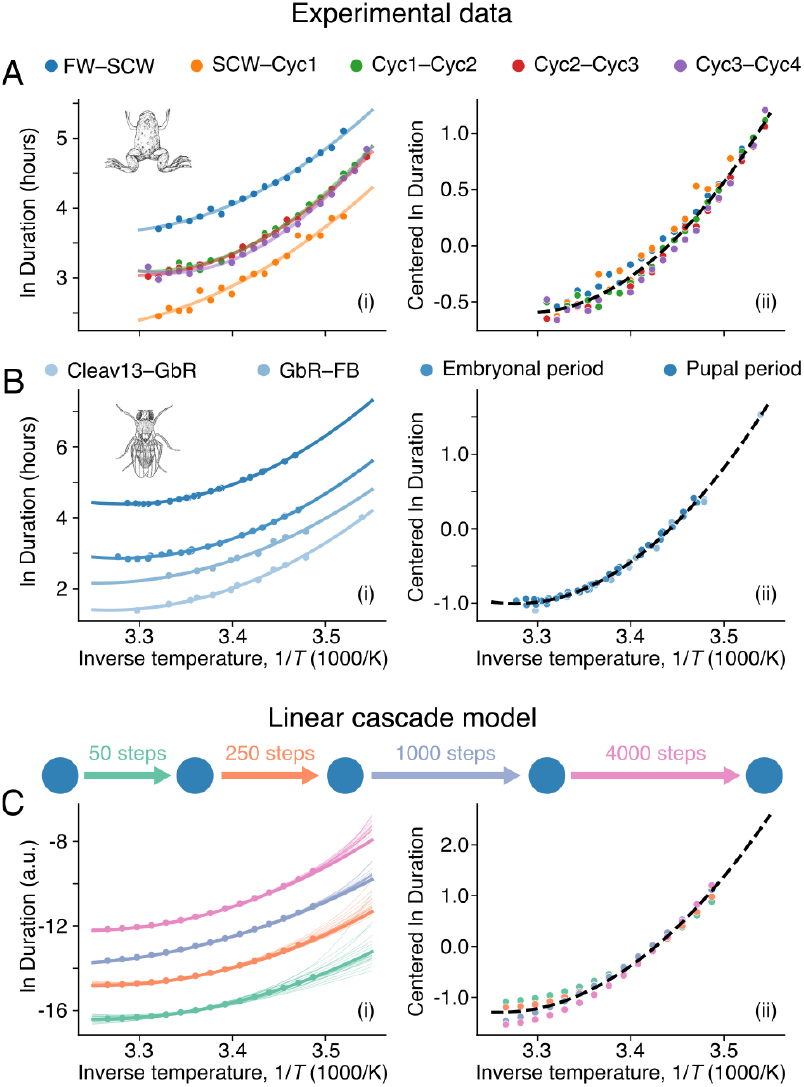
Robustness of biological timing to temperature change. (A) Mean durations of five stages of early embryogenesis in *X. laevis* (Data from Ref. (12)). (B) Mean durations of four developmental stages in *D. melanogaster* (Similar to Figure 5E, data from Refs. (14, 15)). (C) Simulated mean durations of successive developmental stages using linear Markov cascade of increasing size (from 50 to 4000 steps) overlayed with individual realizations. In (A)–(C), left panels display scaling of biological timing per stage, while panels to the right show centered duration, i.e., duration with extracted mean over temperature range per stage, overlapped with the universal scaling curve.

While these examples highlight striking robustness, recent large-scale single-cell profiling in zebrafish shows that this resilience is not absolute: certain cell types are disproportionately sensitive to thermal stress, leading to structural defects (91). Such findings illustrate that robustness at the whole embryo scale can coexist with local vulnerabilities, and that temperature sensitivity can propagate from molecular processes to developmental coordination (92).

The preservation of relative timing despite variable absolute rates is often attributed to active temperature compensation, such as opposing enzymatic reactions with tuned activation energies (93). Our results suggest that it can also arise passively from network structure. In the QE regime, timing emerges from averaging over many similarly scaled, reversible steps, which suppresses variability and preserves stage-to-stage timing ratios even when overall rates change. Simulations (Figure 7C) show that this holds even if stages differ in step number and reaction rates, and remains robust to stochastic variability in individual rates.

At thermal extremes, this robustness breaks down as averaging gives way to a few rate-limiting steps. Variability increases, and relative stage durations can shift, a pattern observed experimentally in some systems at high or low temperatures (93) and consistent with the idea that coordination is maintained only within the Arrhenius-like regime of physiological rates (13). In our framework, such deviations reflect a transition from QE behavior to Arrhenius-like regimes at the edges of the viable temperature range.

#### Temperature compensation

In chronobiology, the maintenance of relative timing across temperature is sometimes called temperature compensation of phase relationships. For example, the neuronal oscillator in the crab *Cancer borealis* preserves phase relationships across temperatures even though its period changes (94, 95). A stricter definition is used for systems such as the circadian clock, where the period itself is temperature-independent (*Q*_10_≈1) (96–99). Mechanistic explanations for strict compensation include antagonistic balance—opposing processes with matched activation energies (100, 101), and enzyme-limited schemes, where several reactions share a common catalytic bottleneck (102, 103). Network-based models have also shown that temperature robustness can emerge from architecture and feedback alone (104–106).

Our framework extends these ideas: compensation can emerge not only from fine-tuned opposing processes but also from distributed averaging across many steps with constrained activation energy distributions. In some cases, this averaging naturally produces similar scaling for opposing reactions, leading to antagonistic balance without explicit tuning. In others, large control networks in the QE regime can exhibit very low apparent activation energies, 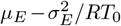, balancing mean and variance in individual reaction activation energies to achieve robustness.

### Negative activation energies

Negative apparent activation energies have long been reported in experimental studies of thermal scaling, particularly at warm temperatures, but their mechanistic basis has remained unclear.

At temperature extremes, they may arise from processes that deviate from simple Arrhenius kinetics, for example, biphasic behavior such as protein denaturation (107, 108), or nonlinear system dynamics approaching bifurcations (12). In Markov networks, cycles and feedforward motifs (Figure 4C–D) can likewise produce negative slopes when transitions become dominated by reverse steps or competing paths. Within the QE regime, negative activation energies can emerge more subtly as apparent network-level properties. Although each microscopic step obeys Arrhenius scaling with a positive energy, motifs such as asymmetric cycles generate local negative slopes, and cascades of such motifs can display an overall negative *µ*_*E*_ (Figure 6). These values therefore do not reflect physically negative microscopic energies, but rather emergent signatures of distributed, multi-step regulation.

Biologically, such mechanisms may explain why negative *µ*_*E*_ values are enriched in organisms adapted to extreme or fluctuating environments (Figure 5D), where they may represent adaptive strategies for maintaining coordination under thermal stress.

### Evolutionary and ecological implications

Across approximately 100 datasets, we find that many biological systems predominantly operate within the QE regime, yet deviations, particularly at elevated temperatures, are frequently observed. This pattern suggests that biological systems may have evolved to function optimally within a buffered thermal window, outside of which developmental timing becomes increasingly erratic or prone to failure. Supporting this idea, recent work (12) showed that, across multiple species, the native environmental temperature closely aligns with the minimum of the QE scaling curve. In other words, these organisms appear to have evolved to maximize developmental speed while remaining within the more robust, low-variability region of the QE regime. These findings carry important implications for understanding how organisms adapt to thermal environments and for predicting the biological consequences of climate change. Consistent with this view, Begasse et al. (13) observed that the fertile temperature window in *C. elegans* and *C. briggsae* overlaps with the range in which cell division adheres to Arrhenius kinetics, highlighting a correlation between developmental robustness and fertility limits. Our results emphasize the importance of characterizing thermal response curves at high resolution, not just at a few “representative” temperatures, to uncover the underlying structure of thermal sensitivity and to identify critical tipping points in developmental systems.

### Connections to empirical biophysical models

Our results also relate to the Sharpe–Schoolfield model, a biophysically inspired framework that combines temperature-dependent reaction kinetics with reversible enzyme inactivation at temperature extremes (23, 24, 109–111). Like our model, it captures three distinct thermal regimes: intermediate speeding up with quadratic-like curvature, a linear scaling at cold extremes, and a slowdown with negative activation energy at hot extremes. However, it attributes the deviations at both ends to enzyme inactivation. In contrast, our Markov framework shows that similar triphasic scaling can emerge from statistical averaging and network effects, even without assuming enzyme denaturation. This provides a mechanistic foundation for empirical patterns observed across taxa. Other commonly used models, such as the QE and DE, typically capture only one or two scaling regimes; while useful for fitting smooth or abrupt deviations, they often fail to describe the full thermal range seen in biological data.

### Toward predictive models of temperature scaling

Together, these insights highlight the need for mechanistic models that explain not only what scaling behavior occurs, but why, and how it might shift under environmental change. By combining statistical physics with biological realism, the Markov cascade model provides a versatile tool for studying temperature-dependent timing. It unifies empirical observations across diverse taxa and stages, and offers mechanistic explanations for scaling deviations that previously lacked theoretical grounding. As global temperatures continue to rise, such generalizable models will be essential for identifying thresholds of developmental failure and for predicting species-specific vulnerabilities.

## Data availability statement

The numerical codes to reproduce the figures in this study are openly available in GitLab (112), and as an archived repository on RDR by KU Leuven [Upcoming]. The experimental datasets used and their analysis is also openly available on RDR [Upcoming].

## ACKNOWLEDGEMENTS

L.G. acknowledges funding by the KU Leuven Research Fund (grant number C14/23/130 and IDN/25/007) and the Research-Foundation Flanders (FWO, grant number G074321N). We thank Stephan Grill for providing us with the data from Ref. (13), and Daniel Cebrián-Lacasa for feedback on the manuscript.

## Supplementary Note 1: The two-ODE embryonic cell cycle oscillator model

The two-ODE cell cycle model is given by the following set of ordinary differential equations

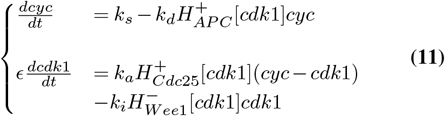

Here, 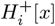 and 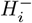 are rising and falling Hill functions, respectively, representing an increase or decrease in the activity of protein *i* in response to changes in *x*:

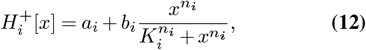

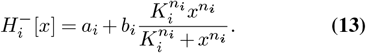

The parameters *a*_*i*_, *b*_*i*_, *n*_*i*_, and *K*_*i*_ denote the basal activity, maximum increase in activity, Hill coefficient, and activation threshold, respectively (12).

### A. Nullclines and dynamics

To analyze the system’s behavior, we examine its nullclines—curves in the (*cyc, cdk1*) phase space along which the time derivative of one variable is zero. The *cyc*-nullcline is defined by setting 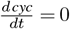,producing a curve in which the synthesis and degradation of *cyc* are balanced. Similarly, the *cdk1*-nullcline is obtained by setting 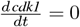, describing where activation and inactivation of *cdk1* are in equilibrium.

The intersection points of the nullclines correspond to steady states of the system. Depending on the location and stability of these points, the system may exhibit monostability, bistability, or oscillatory behavior. In particular, the interaction of fast positive feedback and slower negative feedback can give rise to a stable limit cycle—an isolated, closed trajectory in phase space that corresponds to sustained oscillations in time.

As shown in Figure 2C–D, the concentrations of *cyc* and active *cdk1* oscillate periodically over time (panel C), and the system traces out a loop in the (*cyc, cdk1*) phase plane (panel D). This loop follows a characteristic relaxation oscillator pattern: slow accumulation phases alternate with rapid transitions, producing sharp transitions between activity states (e.g., from low to high *cdk1* activation).

### B. Temperature dependence

The constants *k*_*i*_ are rate constants which are assumed to have an Arrhenius dependence:

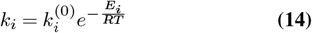

This introduces a temperature dependence in the model and allows to study the temperature dependence of the limit cycle oscillations. An example in which the period has been plotted in function of the temperature is shown Figure 2F (ODE model).

In numerical simulations, it has been observed that when a limit cycle is observed at intermediate temperatures, when *E*_*d*_ ≠ *E*_*s*_ it gets destroyed in the high- or low-temperature limit(12). This can be explained, based on the relative positions of the nullclines in (*Cyc,Cdk1*) phase space. The position of *Cyc*-nullcline is determined by the ratio

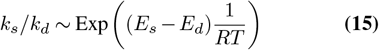

If *E*_*d*_ ≠ *E*_*s*_, then the *Cyc*-nullcline shifts up or down with changing temperature untill it interesects the *cdk1*-nullcline in the upper or lower branc, which creates a fixed point and destroys the limit cycle. This is shown in Figure 2E for the case *E*_*d*_ *> E*_*s*_.

### C. Simulations

Figure 2 in the main text contains a number of simulations of this model and its temperature dependence (C: time series, D: phase space, E: nullclines, F: temperature dependence). For all of these simulations we used the parameters in Table S1 which are taken from (12).

## Supplementary Note 2: Markov jump processes and mean first passage times

In this work, we approximate biological systems as Markov jump processes. Generally speaking, a Markov process is a stochastic process characterized by the key property of memorylessness: the future state of the system depends only on its present state, not on its past history. That is, for a random time-dependent variable *X*(*t*) undergoing a Markov process, the probability distribution of *X*(*t* + *δt*) depends only on the current state at time *t*.

In Markov jump processes specifically, *X*(*t*) can only occupy one of a finite set of discrete states *S* = {1, 2, …, *n*}, while transitions between states occur at random times in continuous time *t*. The likelihood of a transition from state *i* to state *j* is governed by the transition rate *κ*_*ij*_. Informally, if the system is in state *i*, the probability of transitioning to state *j* within a small time interval *δt* is approximately *P* [*i* → *j*] ≈ *κ*_*ij*_*δt*.

Since we are particularly interested in timing, it is useful to define a transition time:

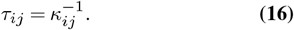

This transition time *τ*_*ij*_ represents the average time to make the transition from state *i* to state *j*, assuming no other transitions from *i* occur during that period.

A Markov process can be conveniently represented as a weighted, directed graph: nodes correspond to discrete states, and directed edges with weights *κ*_*ij*_ represent transitions *i* → *j*^3^. Examples of such graphs are shown in Figures 2–S1. An equivalent representation is the generator matrix *G*, with entries *G*_*ij*_ = *κ*_*ij*_ for *i* ≠ *j* and *G*_*ii*_ = ∑_j≠*i*_ *κ*_*ij*_.

We aim to study the temperature dependence of biological processes. To this end, we postulate that each transition time in the Markov jump process is temperature-activated according to an Arrhenius law:

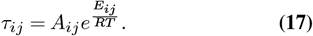

These transitions should be understood as small-scale thermodynamic events that are sufficiently well described by the Arrhenius equation within the relevant temperature range.

Real biological systems are not accurately described by arbitrary state graphs. In regulatory developmental systems in particular, we know they have evolved to reliably reach a specific final state under appropriate conditions (113). We therefore restrict our analysis to simply connected, directed state graphs in which state *n* is the only possible terminal state. In the language of Markov chains, *n* is the unique absorbing state, with total escape rate 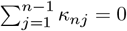. We also assume that the initial state is known and fixed, so that the system always starts in state 1: *X*(*t* = 0) = 1.

In this setting, a natural quantity to study is the time 𝒯 it takes for the process to reach the final state *X* = *n* for the first time, starting from *X* = 1. This is known as the first passage time. Since 𝒯 depends on the stochastic realization of the process, it is a random variable. Markov theory provides tools to compute its statistical moments (18, 43). The most informative of these is the mean first passage (MFP) time, 𝔼 [𝒯], particularly because directed biological systems are expected to minimize higher-order moments through evolution.

The MFP time from state *i* to the absorbing state *n* can be computed by solving the following system of equations:

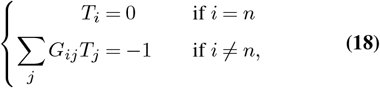

where *T*_*i*_ = 𝔼 [𝒯 |*X*(0) = *i*] is the MFP time from state *i* to *n*, and *G* is the generator matrix of the Markov process.

## Supplementary Note 3: Markov approximation of the two-ODE embryonic cell cycle oscillator

The goal of this supplementary note is to give a detailed overview of how certain features of the two-ODE embryonic cell cycle oscillator can be captured in an approximating Markov jump process. The approximation becomes better as the number of discrete states in the process is increased.

### A. Construction of a four-state Markov chain

To accurately capture the behavior of a temperature-dependent biological system, we need to describe its state and how it evolves over time and with temperature. In the ODE formulation of the embryonic cell cycle model, the state is specified by the continuous concentrations of the proteins Cyc and Cdk1. The system of differential equations in Eq. (12) governs their time evolution, and temperature dependence enters through the rate constants in Equation 14.

We aim to approximate the behavior of this ODE system using a Markov jump process. The first step is to replace the continuous state space with a discrete one. In first instance, we limit *Cyc* and *Cdk1* to two binary states: ON (1) or OFF (0). The ON state represents high Cyclin concentration and high Cdk1 activity, while the OFF state corresponds to low Cyclin concentration and low Cdk1 activity. Combining the states of the two proteins yields a discrete state space with four possible configurations: (0,0), (1,0), (1,1), and (0,1) (see Figure 2B, left).

The next step is to introduce time evolution that mimics the behavior of the ODEs. If the ODE model is in a low-*Cyc* state (0,X), it can transition to a high-*Cyc* state (1,X) via the synthesis rate constant *k*_*s*_. Conversely, it can return from (1,X) to (0,X) due to degradation governed by *k*_*d*_. We therefore introduce two corresponding transition times, *τ*_*s*_ and *τ*_*d*_, to allow switching between low and high *Cyc* states in the discrete-state Markov chain. We note that amount of degradation in the ODE model is regulated by *cdk1* via the Hill function 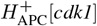.To reflect this, we distinguish between two regimes: 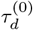 for low *cdk1* ((1,0) → (0,0)), and 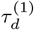 for high *cdk1* ((1,1) → (0,1)). The fast switch from low to high *cdk1* activity ((X,0) → (X,1)) in the ODE model is controlled by the activation term with rate constant *k*_*a*_, which becomes active only when *Cyc* concentration is high. For this reason, we introduce a fast transition time *ϵτ*_*a*_ between (1,0) and (1,1) in the Markov chain. Finally, to complete the model, we add a fast transition time *ϵτ*_*i*_ from (0,1) to (0,0), mimicking *cdk1* inhibition controlled by the rate constant *k*_*i*_ (see again Figure 2B, left).

The final step in constructing the Markov jump process is to introduce temperature dependence. We want the transition times to reflect the same temperature dependence as the rate constants in the ODEs. This can be achieved by assuming an Arrhenius form

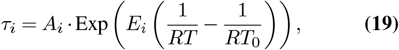

where *E*_*i*_ is taken to be the same as for the corresponding rate constant *k*_*i*_. The prefactors *A*_*i*_ are undetermined at this stage. The only physically motivated constraint is on the ratio between the fast 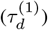 and slow 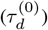 degradation regimes at the reference temperature. This ratio is chosen to match the ratio of minimum to maximum activity of the Hill function

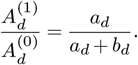

There is no direct relationship between the remaining preexponential factors 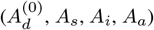 and the rate constants 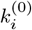. Their numerical values can be determined by fitting to the period of the ODE model (see below).

Once the model is constructed, we extract a period that can be compared to that of the original ODE model. This is done by computing the mean time it takes the Markov chain to return from state (0,0) to (0,0), passing through all other states in the process. Mathematically, this is equivalent to computing the MFP time to (0,0)* starting from (0,0) in a linear cascade of states such as in Figure S1A. From Eqs. (5) and (6), we obtain

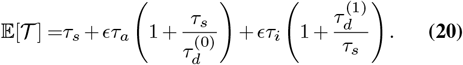

When we fill in all parameters and compare this temperature-dependent MFP time with the period of the ODE model, we find that the ODE model exhibits a stronger temperature dependence (see Figure 2F, *one step* vs. *ODE model*). Specifically, in the Arrhenius plot, the ODE model shows a steeper negative slope in the high-temperature limit and a steeper positive slope in the low-temperature limit. This difference arises because four discrete states are insufficient to fully capture the complex temperature dependence of the continuous ODE model.

### B. A linear cascade of multiple reversible reactions generates a strong temperature dependence

We hypothesize that adding more intermediate states could yield a better approximation of the continuous ODE model. Therefore, in this section, we study in more detail a linear cascade of *n* reversible transitions, with forward transition times *τ*_*f*_, backward transition times *τ*_*b*_, and a final transition time *τ*_0_, as illustrated in Figure S1D. We define the ratio

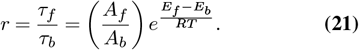

If *E*_*f*_ *< E*_*b*_, this ratio increases in the high-temperature limit; if *E*_*b*_ *< E*_*f*_, it increases in the low-temperature limit.

The MFP time from the initial to the final state is given by

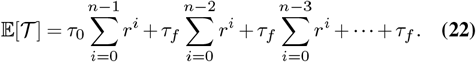

Evaluating the sums yields the closed-form expression:

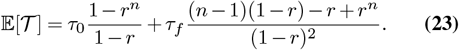

In the limit where *r*≪ 1, backward transitions can be neglected, and the MFP time simplifies to the sum of all forward transition times and the final transition time:

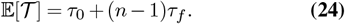

If *r*~ 1, the MFP time begins to increase. Physically, this corresponds to the system becoming trapped in cycles of forward and backward transitions: when *r*~ 1, the system frequently revisits previous states, and as the number of intermediate states grows, the number of potential loops increases, leading to a longer time to reach the end state. When *r*≫ 1, the MFP time is approximately

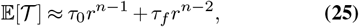

and increases rapidly compared to the *r*≪ 1 case. Note that in this regime, the dependence on *n* becomes exponential.

Since *r* is defined by Eq. (21), it has an exponential dependence on temperature. This implies that the transition from the *r*≪ 1 to the *r*≫ 1 regime occurs abruptly, producing a sharp, temperature-dependent increase in the MFP time in the thermal limits where *r* becomes large. Moreover, this effect is amplified for larger values of *n*.

For the regime *r <* 1, we can also consider the large-*n* limit of Eq. (23). To obtain a finite expression in this limit, we rescale the forward and backward transition times as 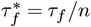 and 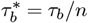. Physically, this reflects the idea that introducing an infinite number of intermediate states only makes sense if the transition times become infinitesimally small. Note that 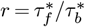. Taking the limit *n* → ∞ while holding 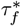 and 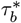 fixed yields:

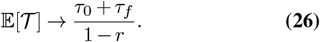

For future reference, we note that this large-*n* limit does not yield a finite expression when *r* ≃ 1.

#### C. Adding extra steps to the Markov chain reproduces the temperature dependence of the ODE model

In the previous section, we showed that adding intermediate states to a linear cascade of reversible processes can produce a steeper temperature dependence of the MFP time in the thermal limits. This motivates us to extend the four-state Markov chain by adding additional steps, such that the MFP time of the chain more closely matches the period of the ODE-based cell cycle oscillator when plotted in an Arrhenius diagram.

In numerical simulations, we observe that Cdk1 activation and inhibition occur rapidly compared to the synthesis and degradation (see Figure 2C). Based on this, we introduce additional steps in the chain, specifically for the synthesis and degradation processes of Cyc. Instead of modeling each process as a single step, we approximate them as *n* discrete reversible steps, each with identical forward and backward transition times. The resulting Markov chain is illustrated in Figure 2B. Again, we can map one period of the chain to the a linear cascade of forward and backward reactions such as in Figure S1B. This chain can be seen as two consecutive cascades as in figure S1D. Therefore its MFP time can be calculated by applying Eq. (5) to each of the individual chains and summing the results. In the continuous limit *n* → ∞, the chain should approximate the dynamics of the original ODE model. From Eq. (26) applied to the two consecutive chains, we find:

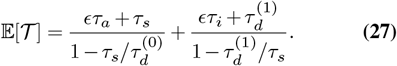

We used this expression to fit the MFP time of the Markov chain to the period of the ODE model as a function of temperature by tuning the unknown pre-exponential constants. A comparison is shown in Figure 2F. Once these constants are determined, we can compare the MFP time for finite values of *n* to the ODE model period. We find that *n* = 50 already gives a close approximation within the temperature range where the ODE model exhibits a limit cycle. Only near the thermal boundaries, where the limit cycle is lost, that is when 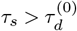 and when 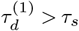 (see Eq. (27)), does the MFP time fail to fully capture the temperature dependence.

This alternative formulation of the cell cycle model as a Markov chain offers new insights into the period increase and eventual breakdown of oscillatory dynamics in thermal limits. The MFP time for Cyc synthesis follows Eq. (5), with

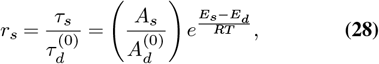

and similarly for degradation

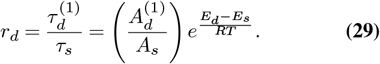

If *E*_*d*_ ≠ *E*_*s*_, both ratios are exponentially temperature-dependent. Within the functional temperature range of the oscillator, both synthesis and degradation must be forward-directed, meaning *r*_*s*_, *r*_*d*_ *<* 1 and

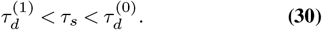

As the temperature changes and *r*_*d*_ → 1, the MFP time of degradation increases dramatically and the total MFP time diverges. Physically, the system becomes trapped in forward-backward loops during degradation, and the limit cycle is destroyed. Likewise, if *r*_*s*_ → 1, backward cycling dominates during synthesis, again disrupting the periodic behavior. Since *r*_*s*_ and *r*_*d*_ have opposite temperature dependencies, one of these breakdowns will occur in the high-temperature limit and the other in the low-temperature limit. This behavior is illustrated in Figure 2G.

## Supplementary Note 4: Temperature scaling of a sum of random transition times: Arrhenius vs. quadratic exponential

In many cases, the MFP time of a biological system can be approximated as a the sum of a large number, *N*, of (random) effective transition times which scale Arrhenius-like around some reference temperature *T*_0_. Examples given in this paper include Eq. (5) and Eq. (42). In this section we derive the temperature scaling of such a system and show that two possible forms of scaling emerge at different temperature ranges: QE scaling when we can average over a large number of effective transition times and Arrhenius scaling when one of the transition times dominates.

## A. Distribution from the central limit theorem

Assuming forward propagation *r*≪ 1 in the cascade and that preexponential factors *A*_*i*_ and activation energies of forward reactions *E*_*i*_ are random variables drawn from arbitrary distributions, we can rewrite Eq. (5) as

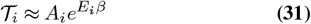

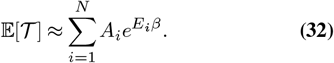

In this expression, we took *β* = 1*/*(*RT*) −1*/*(*RT*_0_) such that *A*_*i*_ is the effective transition time at the reference temperature. In the limit *N* → ∞, the central limit theorem assures that if all exponentials are independent and identically distributed random variables, then 𝔼 [𝒯] will be normally distributed with mean *µ* and variance *σ*^2^ over different realizations, where *E*_*i*_ and *A*_*i*_ (*i* = 1,.., *N*) are randomly drawn from their respective distributions

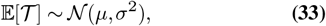

where

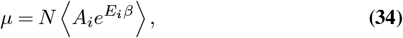

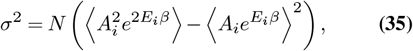

where ⟨…⟩ denotes the mean with respect to the distributions of *A*_*i*_ and *E*_*i*_ (*i* = 1,.., *N*).

Rewriting Eq. (34) as 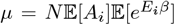,we express 𝔼 [*A*_*i*_] = *µ*_*A*_ and use the cumulant form for 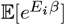:

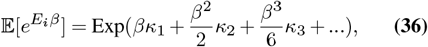

where *κ*_1_ = *µ*_*E*_ and *κ*_2_ = Var[*E*]. Neglecting cumulants of order 3 and higher, one gets a closed-form expression for the MFP time:

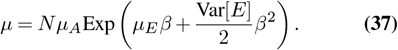

Note that this general form is valid for arbitrary distributions of *A*_*i*_ and *E*_*i*_.

Likewise, we find the variance of MFP time:

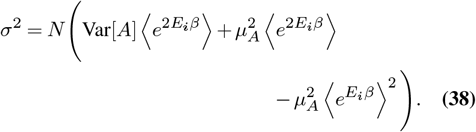

At extremal temperatures the variance in *A*_*i*_ at the reference temperature can usually be neglected which means that

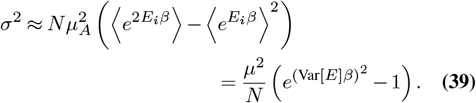

### B. Relative variation

We note that *µ* from Eq. (37) has a QE temperature dependence. Whether 𝔼 [𝒯] will follow this temperature scaling of *µ* for a typical realization drawn from the distributions of *A*_*i*_ and *E*_*i*_, depends on the relative variation

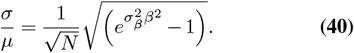

In the region where 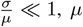 is a good approximation of 𝔼 [𝒯] for all realizations, i.e., simulations of a system each with a given set of (*A*_*i*_, *E*_*i*_), thus the system scales according to a QE law. If 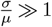, which is the case for both hot and cold temperature extremes, energy variation Var 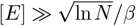 contributes to a 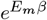 of the MFP time. The picture that emerges is that of two different regimes with a boundary at extremal temperatures that is located at *σ/µ* = 1 or at

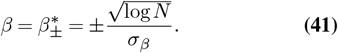

## Supplementary Note 5: Linear cascades of reversible processes with random transition times

In this section, we approximate biological systems as linear cascades of reversible processes with *n* distinct transition steps, as schematically illustrated in Figure S1E.

### A. The mean first passage time

The MFP time for such a cascade is given by

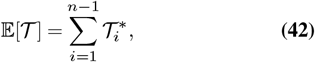

where the 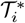 are the effective forward transition times, defined as

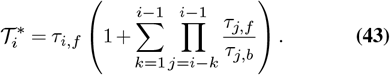

Each 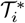 represents the average time required to move from state 1 to state *i* accounting for all possible backward excursions along the cascade, which we refer to as an effective passage time. These effective times reduce the reversible cascade to an equivalent process with purely forward transitions.

### B. Forwards and backwards regime

By analogy with Eq. (21), we define ⟨*r*⟩ = ⟨*τ*_*i,f*_ */τ*_*i,b*_⟩. If ⟨*r*⟩ ≪ 1, it suggests that that on average forward transitions dominate the propagation in the cascade, so we can approximate 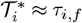 and from Eq. (42) we find

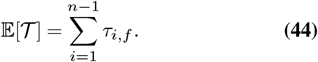

In this forwards regime, we can ignore the backwards transitions in the computation of the MFP time and the system behaves as if there are only forward transitions. If⟨ *r*⟩~ 1 the backward transitions will increase the MFP time. Physically, this happens because the system can get trapped in backward cycles before it exits towards the final state in the forward direction. If ⟨*r*⟩≫ 1, we say we are in the backwards regime. The boundary between the two regimes is determined by the temperature at which ⟨*r*⟩ = 1.

### C. Introducing temperature dependence

Connecting the results in the previous subsections with the rest of the paper, we again assume all transition times have an Arrhenius dependence on temperature

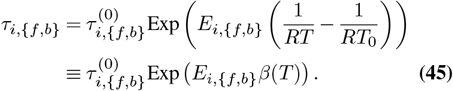

In this, *T*_0_ is a reference temperature at which we suppose the cascade is working in the forward regime and *β*(*T*) = 1*/*(*RT*) −1*/*(*RT*_0_) is introduced to facilitate calculations. We now choose a distribution of transition times by imposing a normal probability distribution 𝒩 (*µ, σ*) with mean *µ* and standard deviation *σ*^2^ on the activation energies and the the transition times at the reference temperature:

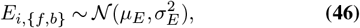

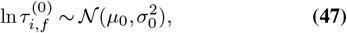

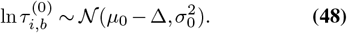

Note that all activation energies are assumed to be identically distributed, but the forward and the backward transition times have a different mean at the reference temperature which is controlled by 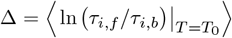. In the forward working regime we expect that Δ *<* 0.

### D. Temperature-dependent distributions of forward and backward transition times

Assuming that the activation energies as well as the logarithms of forward and backward transition times at the reference temperature *T*_0_ are drawn from the normal distributions, Eqs. (46)-(48), distri-bution of *τ*_{*f,b*}_ at temperature *T* is given by:

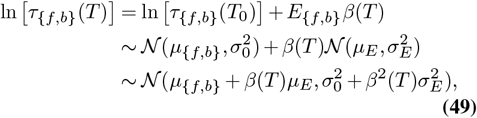

where *µ*_*f*_ = *µ*_0_ and *µ*_*b*_ = *µ*_0_− Δ are means of the forward and backward transition times at *T*_0_. Next, one can derive the distribution of the forward-to-backrward ratio *r* = *τ*_*f*_ */τ*_*b*_ at a given temperature *T* as:

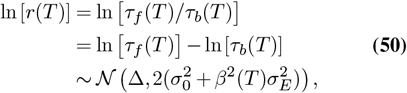

Or, explicitly:

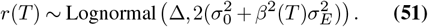

From Eqs. (50) and (51), it follows that only the variance of the forward-to-backward ratio distribution is temperature-dependent, and, importantly, it increases symmetrically relative to the reference temperature *T*_0_ (Figure 3D). In other words, as the system moves away from the reference temperature, the probability of getting *r*_*i*_ *>* 1 increases on both cold and warm sides, amplifying the cycle motifs in the linear transition cascade.

### E. Temperature-dependent distributions of transition times in different populations

Moving forward, we want to get more insight into the development of a heavy tail of the distribution of *r* away from *T*_0_. We take a closer look on the distribution of transition times depending on the balance between respective forward and backward activation energy. Taking into account that in the populations with *E*_*i,f*_ *> E*_*i,b*_ and with *E*_*i,f*_ *< E*_*i,b*_ transition times scale differently, i.e., backward transitions dominate at low temperatures in the former case and at the higher in the later case (Figure 3C), let us treat these two populations separately and find the distributions of conditioned variables *τ*_{*f,b*}_|*E*_*f*_ *> E*_*b*_ and *τ*_{*f,b*}_|*E*_*f*_ *< E*_*b*_.

This problem largely comes down to deriving conditional distributions of respective activation energies: *E*_{*f,b*}_| *E*_*f*_ *> E*_*b*_ and *E* _{*f,b*}_ |*E*_*f*_ *< E*_*b*_ (Figure. S2A). To do so, consider a bivariate distribution (*E*_*f*_, *E*_*b*_), where each component individually is drawn from the normal distribution in Eq. (46). Next, define a vector:

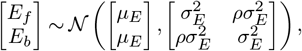

and set *ρ* = 0, assuming no correlation between *E*_*f*_ and *E*_*b*_. To facilitate the analysis, introduce new variables:

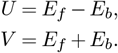

New variables (*U, V*) are obtained by linear transformation (rotation) of (*E*_*f*_, *E*_*b*_), and, thus, are also drawn from a bivariate normal distribution. From the latter it follows that:

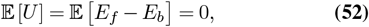

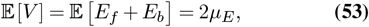

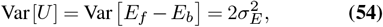

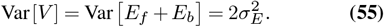

Now, without the loss of generality, consider the distribution of *E*_*f*_ conditioned by *E*_*f*_ *> E*_*b*_, or, equivalently, by *U >* 0. Defining *E*_*f*_ through the new variables as *E*_*f*_ = (*U* + *V*)*/*2, we express sought conditioned variable as:

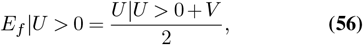

where *U* |*U >* 0 is drawn from a truncated normal distribution with mean and variance:

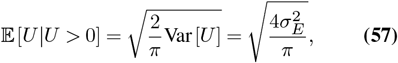

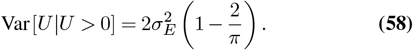

Substituting Eqs. (53), (55), (57), and (58) in Eq. (56), we obtain the distribution of forward activation energies *E*_*f*_ in the population of transitions with *E*_*f*_ *> E*_*b*_:

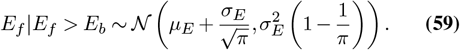

Repeating the same for *E*_*b*_| *E*_*f*_ *> E*_*b*_ and for the *E*_*f*_ *< E*_*b*_ population, one obtains:

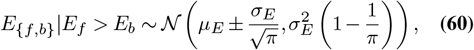

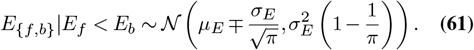

Finally, with known expressions for *E*_{*f,b*}_ |*E*_*f*_ *> E*_*b*_ and *E* _{*f,b*}_ *E*_*f*_ *< E*_*b*_, we find that sought transition times are also normally distributed:

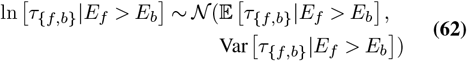

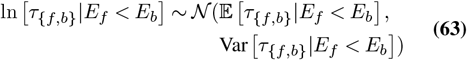

with respective means and variances:

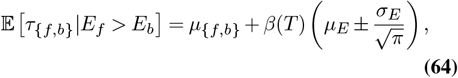

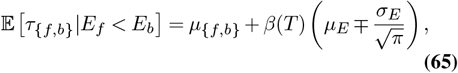

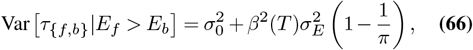

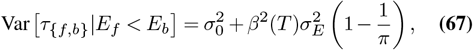

From the above equations, it follows that the variation of transition times have the same temperature dependence in both populations, while the means scale differently: the forward and backward transition times get closer at lower temperatures and diverge at higher temperatures in the *E*_*f*_ *> E*_*b*_ population and the opposite occurs in the *E*_*f*_ *< E*_*b*_ population. In summary, at colder temperatures, *E*_*f*_ *> E*_*b*_ transitions amplify the cycles, while *E*_*f*_ *< E*_*b*_ transitions contribute to the cycles at warmer temperatures (Figure S2B).

### F. Temperature scaling

Let us now take a look at the scaling of the MFP time. Based on the analysis of the forward and backward transition times in the previous section we distinguish between a forward regime where ⟨*r*(*T*) ⟩ ≪1 and the MFP time is dominated by forward transitions, and a backward regime where ⟨*r*(*T*) ⟩ ≫1 and the MFP time is dominated by the effect of backward transitions. Since for the lognormal distribution in Eq. (51) we have

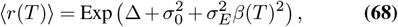

we find that the boundaries between the forward and the backward regime are located at

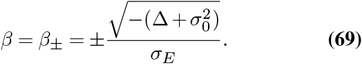

Since the temperature dependence is exponential, the transition between the forwards and backwards regime will happen over a narrow temperature range.

#### F.1. Temperature scaling in the forwards regime

In S4 we derive the temperature scaling of a system which has a MFP time that can be written as the sum of exponentials such as in Eq. (44). The conclusion is that when the number of exponentials is large and *σ*_*β*_ is small, the MFP time scales according to a QE temperature law that emerges from an averaging over all exponentials. Using the results in Eq. (34) and Eq. (41) and applying them to the forwards approximation in Eq. (44) where *σ*_*β*_ = *σ*_*E*_ and *N* = *n*, we find that in the forwards regime

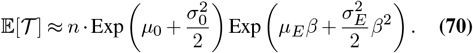

for all temperatures *β* for which

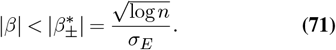

At more extreme temperatures, individual transition times tend to dominate the sum in Eq. (44) and the system will scale Arrhenius-like, following the dominant forward transition time. However, if *n* is sufficiently large enough then 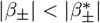 and the MFP time scales quadratically within the entire forward regime.

#### F.2. Temperature scaling in the backwards regime

To understand what the temperature dependence of the MFP time in the temperature region ⟨*r*⟩ *>* 1 looks like, we can start from Eq. (43) and note that with Eq. (45) the effective transition times take again the form of a sum of exponentials

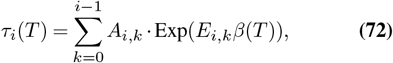

with

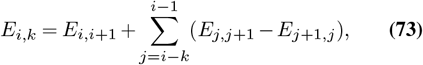

and *A*_*i,k*_ a constant factor that is determined as the product and quotient of pre-exponential factors of individual transition times. The random activation energies *E*_*i,k*_ can be positive or negative. The variance on these effective activation energies is much higher than that on the individual forward activation energies since

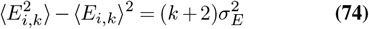

This means that 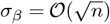 while the number of effective transition times is

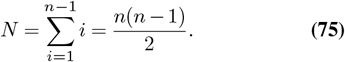

We conclude that

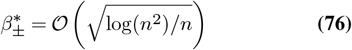

and that 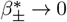 if *n* → ∞. Thus, in the large-number limit, there will be no region of QE scaling where we can average over a large number of transition times and the MFP time always scales Arrhenius-like in the backwards regime.

The exponential with 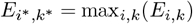 will dominate the MFP time in the high-*β* or low-temperature limit and cause a positive linear slope in the Arrhenius diagram. The exponential with 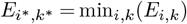 will dominate in the high-temperature limit and will typically cause a negative linear slope in the Arrhenius diagram.

### G. Numerical simulations

To check these theoretical results we conducted a series of numerical simulations of linear cascades with transition times drawn from a random distribution.

For a first round of simulations we used the parameter values in Table S2 (middle column) and *n* variable and used these parameters to assign temperature-dependent random transition times to each of the transitions in the linear cascade via Eqs. (46), (Eq. (47)), and (48). The motivation for the parameters of the activation energies comes from (44) which gives a survey of a large number of enzymatic reactions and their temperature dependence. Afterwards we calculated the MFP time of this cascade in function of temperature via Eq. (42) for a temperature range of 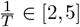 · 1000/*K*. In the QE temperature region we compared the MFP time with the theoretical prediction Eq. (70). Outside of this region we determined a best fit of a linear Arrhenius function with positive activation energy in the low-temperature and negative activation energy in the high-temperature region to the MFP time. This process was repeated 100 times for different values of *n*. The results are given and discussed in Figure 3F and the accompanying text. We also repeated this procedure for a uniform distribution of activation energies with the same mean and standard deviation as the normal distribution and reported no significant differences, confirming the reliability of the approximation via a normal distribution.

In a second round of simulations we repeated the same procedure but now in a setup with a feedforward loop. The exact parameters that were used are given in Table S2 (right column). The network topology is the same as in the first row but contains an additional shortcut as shown in Figure 3B, now contains a feedforward loop with fixed transition times that has been engineered to become active at low temperatures. We did this by making *E*_control_ ≪*µ*_*E*_ such that at low temperatures *τ*_control_ ≪*τ*_*i,f*_ for an average *τ*_*i,f*_. Therefore, the shortcutted path becomes much more probable at low temperatures and will always be taken. The results for the simulations are given in Figure 3E and G and discussed in the accompanying text.

## Supplementary Note 6: Linear cascade of backward cycles with random transition times

In this section we derive the temperature scaling of the network topology in Fig. 6D and show how it can lead to negative QE temperature scaling in a certain temperature range.

### A. The mean first passage time

The MFP time is the sum of the MFP time of *n* − 1 individual cycles (see Eq. (92))

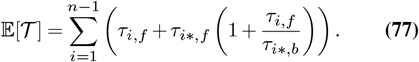

### B. Temperature scaling

As before we call 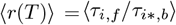,We can again distinguish between a forward regime when ⟨*r*⟩ ≫ 1 and a backwards regime when ⟨*r*⟩ ≪ 1.

#### B.1. Forward regime

In the forward regime, the contribution from the backward cycles to the temperature scaling can be neglected. The MFP time is the sum of 2(*n*− 1) forward transition times. We assume a normal distribution 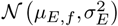 for the forward activation energies and a mean of *µ*_0,*f*_ for the forward pre-exponential factors. From Eq. (34) we find that in the large number limit

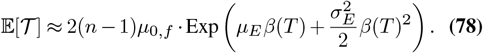

For finite-sized systems we expect to see deviations from the QE in the form of Arrhenius scaling at cold temperatures corresponding to

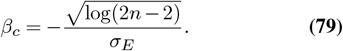

#### B.2. Backwards regime

In the backward regime the MFP time will be dominated by the contributions from the backward cycles and we can neglect the individual forward reactions in Eq. (77) to find

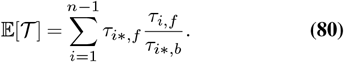

We assume a normal distribution 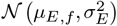 and 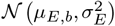 for the forward and backward activation energies and a means *µ*_0,*f*_ and *µ*_0,*b*_ for the forward and bakcward pre-exponential factors. Again using, Eq. (34) we find that in the large number limit a QE scaling law emerges

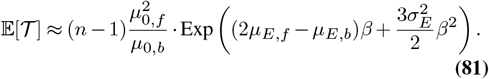

If *µ*_*E,b*_ *>* 2*µ*_*E,f*_ then this will be a negative QE. For finite-sized systems we expect to see deviations from the QE in the form of Arrhenius scaling at hot temperatures corresponding to

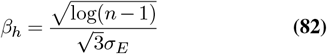

Typically, it will be the individual exponentials with large negative activation energies that will dominate the MFP time at these extremal temperatures.

### C. Total temperature scaling

Suppose that we are dealing with a finite-sized system, that *r*(*β*^*^) = 1 and that *µ*_*E,b*_ *>* 2*µ*_*E,f*_. Based on the discussion above we can then discertain between four different regions of temperature scaling

1. Arrhenius scaling at the hot side when

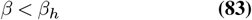
2. Negative QE scaling at the hot side with a large curvature when

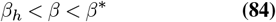
3. QE scaling with a smaller curvature at the cold side when

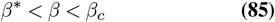
4. Arrhenius scaling at the cold side when

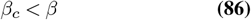

### D. Generalizations

In the examplatory topology of this section, the difference between the forward and backward activation energies should quite high to get a negative QE in the bakcwards regime. However, one could also consider cascades of cycles with *m* forward and backward reactions where *m >* 1. In the backwards regime, the largest exponentials would dominate and the approximate MFP time would become

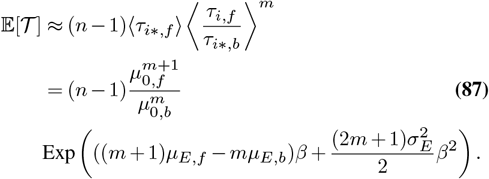

In this case one would get a negative QE for 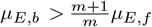 which makes that the difference between the forward and backward activation energies need not be large to get a negative slope. Note that the curvature also increases for larger backward cycles.

If multiple backward cycles with different lengths would be present in the same cascade, the cycles with the longest length would always dominate in the backward regime and the other ones could be ignored.

## Supplementary Note 7: Simple motifs with a negative activation energy

One of the main paradoxes to be solved, is how the overall negative activation energy emerges from a series of small-scale transitions which all have positive activation energies. In this section, we discuss one example that illustrates when this is not possible, and two minimal examples of state graph topologies for which this is possible. The two last motifs highlight the two physically different mechanisms that might lead to an overall macroscopic negative activation energy in the temperature scaling of the MFP time.

### A. Cascade of sequential forward reactions

Consider a simple cascade of forward-directed transitions such as depicted in Figure S1C. The MFP time is just the sum of all individual transition times

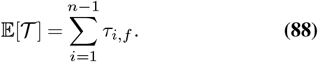

If we calculate the temperature dependence of 𝔼 [𝒯] we find that

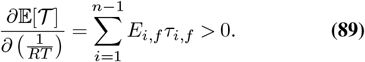

We conclude that the MFP time will never show a negative slope in the Arrhenius diagram if there is only one possible path towards the final state. An example with two forward transitions is shown in the bottom half of Figure 4B. To make this plot we used the parameter in Table S3 (left).

### B. A cycle

Consider the simplest state graph with one cycle such as depicted in Figure 4D 4C??. Calculating the MFP time, using Equation Eq. (42) and Eq. (43), results in

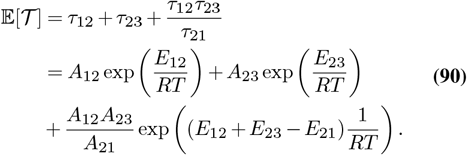

Besides the sum of the two forward transition times, an extra term appears. It captures the possibility of trapping the system in the cycle (1) ↔(2) a number of times before it exits towards state (3).

Look at the particular case: *E*_21_ *> E*_12_ + *E*_23_. In the lowtemperature limit 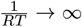, systems scales Arrheniuslike with a positive activation energy *E* = max{*E*_12_, *E*_23_}. But when 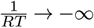 the following scaling is observed

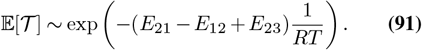

However,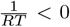 does not correspond to physical temperatures. Therefore, it depends on the pre-exponential constants whether this limit is already observed for physical temperatures with *T >* 0. If this is the case, 𝔼 [𝒯] scales Arrhenius-like with a negative activation energy in the high-temperature limit.

An example of this scaling behaviour can be seen in the bottom half of Figure 4C. To make this plot we used the parameter in Table S3 (middle).

### C. A feedforward loop

Consider a minimal feedforward loop without cycles but with two forward directed paths such as in Figure 4E. The MFP time is given by

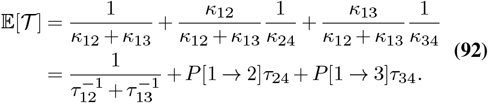

Where *P* [1 → 2] is the chance that the path along state (2) is taken and *P* [1 → 3] is the chance that the path along state (3) is taken.

Take *E*_12_ *< E*_13_ and *τ*_12_, *τ*_13_ ≪*τ*_24_, *τ*_34_. This setup makes that the followed path, is controlled by the first transition step, but the total MFP time is entirely dominated by the second step. In the low-temperature limit *τ*_12_ ≪*τ*_13_. Therefore the path along state (2) is taken and 𝔼 [𝒯] ≈*τ*_24_. In the high-temperature limit, the reverse could happen which would result in 𝔼 [𝒯] ≈ *τ*_34_.

Note that in both of these limits the local activation energy is constant and positive. However, if the transition between these two limits happens fast and *τ*_34_ ≪*τ*_24_ the local activation energies in the transition between the two limits can be negative. An example for which this happens is shown in the bottom half of Figure 4D. To make this plot we used the parameters in Table S3 (right).

## Supplementary Note 8: Data analysis

### A. Dataset construction

We conducted a literature search to find studies measuring the timing of various biological processes under temperature variation. We included recordings of durations of early and later embryonic development stages in different model organisms, as well as bacteria and phytoplankton growth rates. When available, we extracted data from public online repositories or article tables; otherwise, we used `PlotDigitizer` (https://plotdigitizer.com) to get data from the article figures. Overall, we collected 121 datasets from 48 source, resulting in a total amount of 2174 data points (18 points per set on average, minimum of 5 points per set). On average, temperature range spanned 30±17 *K* with the minimum of 10 *K*.

### B. Fitting procedure

Fitting procedure aimed to select and fit the best model explaining temperature scaling of biological timing for individual curves. The candidate models were:

1. **Quadratic-exponential (QE)**, which supposedly describe scaling of biological timing in a wide range of temperatures following Markov-chain formalism (See Eq. (7)):

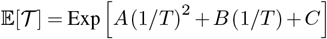

with 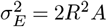 and 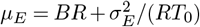;
2. **Quadratic-linear-exponential (QLE)**, capturing deviation of biological timing from the QE model at *either* cold *or* hot temperature extreme:

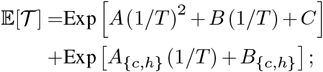
3. **Quadratic-double-linear-exponential (QDLE)**, capturing deviation of biological timing from the QE model at *both* cold *and* hot temperature extremes:

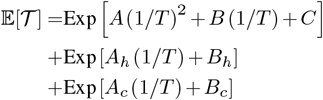

In both QLE and QDLE models, *E*_{*c,h*}_ = *RA*_{*c,h*}_.

To find and fit the best model to the data, we used a multi-step fitting procedure described below.

- **Step 1:** quadratic function was fitted on the plane (1000*/T*, ln *τ* (*T*)) in a whole temperature range to the all datasets.
- **Step 2:** datasets were labeled as **QE** based on the goodness-of-fit evaluation (*R*^2^ *>* 0.95) and additional visual inspection for less clean data.
- **Step 3:** datasets, exhibiting deviations from the QE function, were subjected to automated model fitting/selection process. Deviations were identified by linear regression on residuals of a quadratic fit (20% of datapoints at each side). If linear regression coefficient exceeded a predefined threshold, the deviation was identified. Based on the identified deviation (on the hot side, on the warm side, or both) either one-sided **QLE** or two-sided **QDLE** function was fitted.
- **Step 4:** similar to **Step 2**, datasets were labeled as QLE or QDLE based on the goodness-of-fit evaluation (*R*^2^ *>* 0.95) and additional visual inspection for less clean data.
- **Step 5:** datasets, failing the automated fitting, were subjected to a similar model/fitting selection but with the manual setting of deviation boundaries.

Parameter fitting was performed using the least squares algorithm implemented in the `least_squares` function from `scipy.optimize` package for Python. Goodness-of-fit was evaluated using the `r2_score` function from `sklearn.metrics` package for Python. Fitting code was optimized using Generative AI (ChatGPT-5o).

## Supplemental Tables

**Table S1.** All parameters which were used to simulate the 2 ODE cell-cycle model in Figure 2C,D,E,F. All parameters in the upper half are taken from (12) where they are shown to represent aspects of a realistic cell division cycle. The parameters 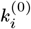 are the values of the rate constants *k*_*i*_ at *T* = *T*_0_. The activation energies in the bottom half are chosen by us to highlight specific aspects of the model.

| Parameter | Value |
| --- | --- |
| $k_s^{(0)}$ | 1.25 nM/min |
| $k_d^{(0)}$ | $0.1 \text{ min}^{-1}$ |
| $k_i^{(0)}$ | $1 \text{ min}^{-1}$ |
| $k_a^{(0)}$ | $1 \text{ min}^{-1}$ |
| $a_{Cdc25}$ | 0.2 |
| $b_{Cdc25}$ | 0.8 |
| $K_{Cdc25}$ | 30 nM |
| $n_{Cdc25}$ | 10 |
| $a_{Wee1}$ | 0.1 |
| $b_{Wee1}$ | 0.4 |
| $K_{Wee1}$ | 30 nM |
| $n_{Wee1}$ | 5 |
| $a_{APC}$ | 0.1 |
| $b_{APC}$ | 0.9 |
| $n_{APC}$ | 15 |
| $K_{APC}$ | 30 nM |
| $\epsilon$ | 0.1 |
| $1/T_0$ | $3.4 \cdot 1000/\text{K}$ |
| $E_s$ | 100 kJ/mol |
| $E_d$ | 60 kJ/mol |
| $E_i$ | 75 kJ/mol |
| $E_a$ | 75 kJ/mol |

**Table S2.** All parameters which were used to simulate the linear cascade of reversible processes in Figure 3. The left column corresponds to panels D and F, the right column to panels E and G.

| Parameter | Reversible transitions | Reversible transitions + feedforward |
| --- | --- | --- |
| $\mu_E$ | 50 kJ/mol | 50 kJ/mol |
| $\sigma_E$ | 10 kJ | 10 kJ |
| $\mu_0$ | -20 | -20 |
| $\Delta$ | -2 | -2 |
| $\sigma_0$ | 0.2 | 0.2 |
| $1/T_0$ | 100/K | 100/K |
| $E_{\text{control}}$ | - | 500 kJ |
| $E_{\text{shortcut}}$ | - | 10 kJ |
| $\log_{10}(A_{\text{control}})$ | 0 | - 20 |
| $\log(A_{\text{shortcut}})$ | 0 | - 20 |
| $n$ | variable | variable |

**Table S3.** All parameters which were used to simulate the three examples in Figure 4. The left, middle and right column correspond to panels B, C and D.

| Parameter | Two sequential reactions | A cycle | Feedforward |
| --- | --- | --- | --- |
| $A_{12}$ | 5 | 0.1 | 0.001 |
| $A_{23}$ | 30 | 0.1 | |
| $A_{21}$ | | 0.1 | |
| $A_{13}$ | | | 0.001 |
| $A_{34}$ | | | 10 |
| $A_{24}$ | | | 0.1 |
| $E_{12}$ | 5 kJ/mol | 30 kJ/mol | 10 kJ/mol |
| $E_{23}$ | 30 kJ/mol | 30 kJ/mol | |
| $E_{21}$ | | 70 kJ/mol | |
| $E_{13}$ | | | 100 kJ/mol |
| $E_{34}$ | | | 10 kJ/mol |
| $E_{24}$ | | | 10 kJ/mol |

## Supplemental Figures

**Fig. S1.**
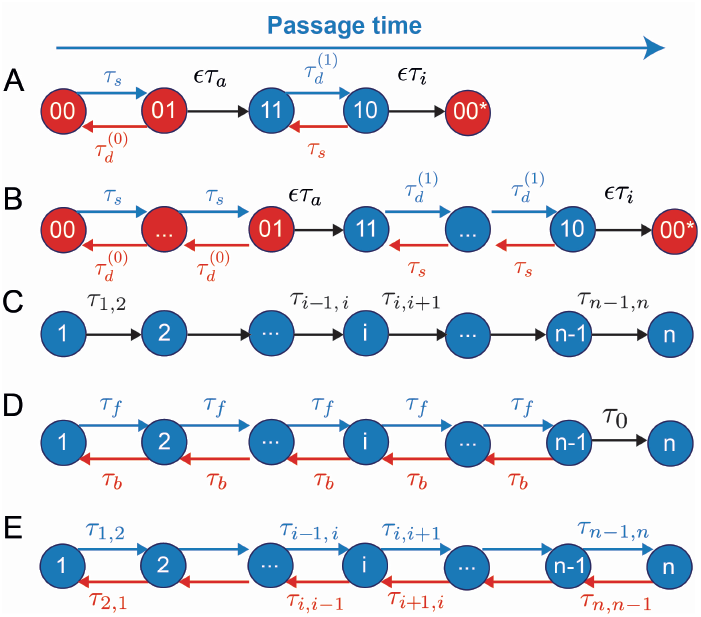
Different linear cascades of reactions which were considered to explain the temperature dependence of biological networks. (A) One period of the minimal Markov approximation of the two-ODE cell cycle oscillator mapped to a linear cascade (Section A) (B) One period of the Markov approximation of the two-ODE cell cycle oscillator with additional states (Section C). (C) Cascade of forward reactions. This cascade has no macroscopic negative activation energy (Section B). (D) Cascade of fixed reversible reactions followed by a forward reaction. This cascade can produce a strong temperature dependence with negative activation energies in the high-temperature limit (Section C). (E) This cascade is the most general one and can produce negative activation energies in the high-temperature limit and a QE scaling at intermediate temperatures (section A)

**Fig. S2.**
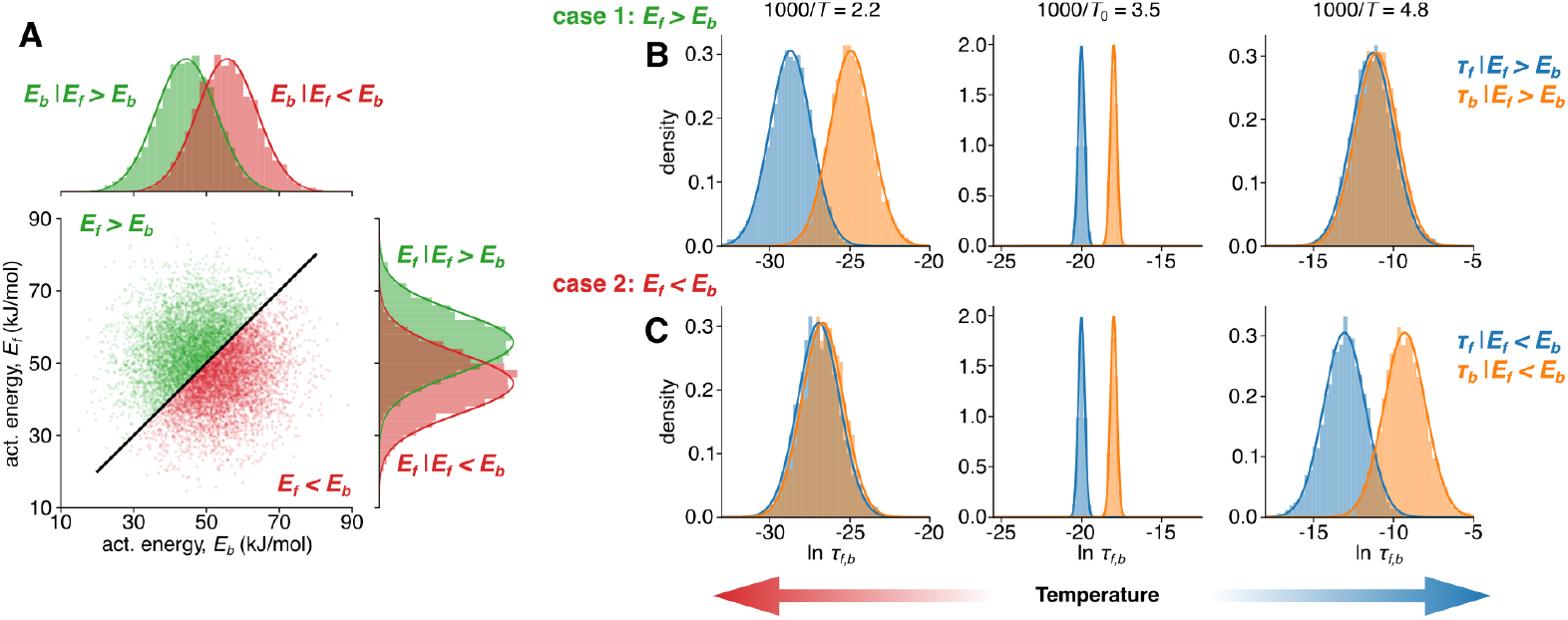
Temperature-dependent distributions of forward and backward transition times. (A) Scatterplot showing bivariate distribution of *E*_*f*_ versus *E*_*b*_. Histograms to the top and to the right show distributions of *E*_*f*_ and *E*_*b*_ conditioned by *E*_*f*_ *> E*_*b*_ (green) and *E*_*f*_ *< E*_*b*_ (red). (B) Distributions of transition times *τ*_*f*_ (blue) and *τ*_*b*_ (orange) conditioned by *E*_*f*_ *> E*_*b*_ (top row) and *E*_*f*_ *< E*_*b*_ (bottom row) at different temperatures.

**Fig. S3.**
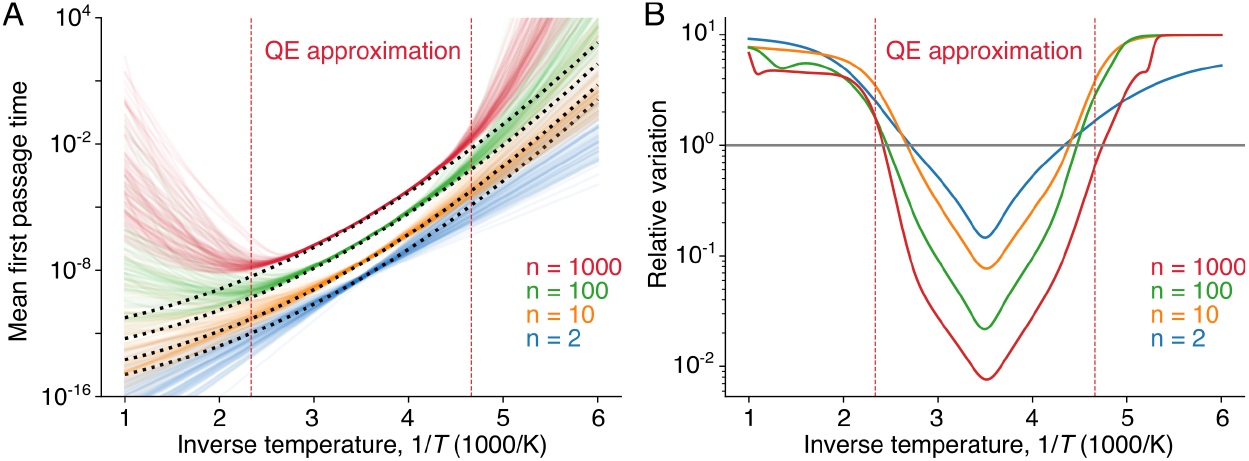
The mean first passage time varies little in the intermediate temperature range. Simulation of the temperature dependence of the MFP time for system (A) and in Fig. 3. Individual transition times are drawn from a distribution with *µ*_*E*_ = 50 kJ/mol, *σ*_*E*_ = 10 kJ/mol, *µ*_0_ = −20, *σ*_0_ = 0.2, *µ*_*r*_ = 2. In the intermediate temperature regime, the QE prediction from Equation (7) provides a good fit. Boundaries of the QE regime are determined from Equation (**??**). (A) The MFP time as a function of temperature for 100 realizations of transition rate distributions. (B) The relative variation (standard deviation/mean) of MFP time across realizations as a function of temperature.

## Footnotes

1 Here *Q*_10_(*T*) ≡ *κ*(*T* + 10^°^C)*/κ*(*T*) is the *local* fold-change in rate per 10^°^C. For Arrhenius *κ*(*T*) = *A* exp(−*E*_*a*_*/RT*), 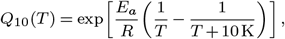 so *Q*_10_ depends on both *T* and *E*_*a*_ (e.g., at 20^°^C, *E*_*a*_ = 50–80 kJ mol^−1^ gives *Q*_10_≈ 2.0–2.8). For durations *τ* = 1*/κ*, the corresponding factor is 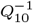. In non-Arrhenius (curved) regimes, a single *Q*_10_ is only a local approximation (12); for a general Δ*T*, the factor is 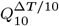.

2 Irreversible forward transitions can also be incorporated by setting the corresponding backward transition time to infinity.

3 Note that edges are often labeled by their transition time *τ*_*ij*_. This implicitly refers to a transition rate of 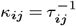.

